# Androgen deprivation-mediated activation of AKT is enhanced in prostate cancer with *TMPRSS2:ERG* fusion

**DOI:** 10.1101/2024.08.26.609679

**Authors:** Fen Ma, Sen Chen, Luigi Cecchi, Betul Ersoy-Fazlioglu, Joshua W. Russo, Seiji Arai, Seifeldin Awad, Carla Calagua, Fang Xie, Larysa Poluben, Olga Voznesensky, Anson T. Ku, Fatima Karzai, Changmeng Cai, David J. Einstein, Huihui Ye, Xin Yuan, Alex Toker, Mary-Ellen Taplin, Adam G. Sowalsky, Steven P. Balk

**Author notes:** Address correspondence to Steven P. Balk, Beth Israel Deaconess Medical Center, 330 Brookline Avenue, Boston, MA 02115. co-first authors (Fen Ma, Sen Chen, Luigi Cecchi).

## Abstract

*TMPRSS2:ERG* gene fusion (*T:E* fusion) in prostate adenocarcinoma (PCa) puts ERG under androgen receptor (AR) regulated *TMPRSS2* expression. *T:E* fusion is associated with *PTEN* loss, and is highly associated with decreased *INPP4B* expression, which together may compensate for ERG-mediated suppression of AKT signaling. We confirmed in PCa cells and a mouse PCa model that ERG suppresses *IRS2* and AKT activation. In contrast, ERG downregulation did not increase INPP4B, suggesting its decrease is indirect and reflects selective pressure to suppress INPP4B function. Notably, INPP4B expression is decreased in *PTEN*-intact and *PTEN*-deficient *T:E* fusion tumors, suggesting selection for a nonredundant function. As ERG in *T:E* fusion tumors is AR regulated, we further assessed whether AR inhibition increases AKT activity in *T:E* fusion tumors. A *T:E* fusion positive PDX had increased AKT activity in vivo and response to AKT inhibition in vitro after androgen deprivation. Moreover, two clinical trials of neoadjuvant AR inhibition prior to radical prostatectomy showed greater increases in AKT activation in the *T:E* fusion positive versus negative tumors. These findings indicate that AKT activation may mitigate the efficacy of AR targeted therapy in *T:E* fusion PCa, and that these patients may most benefit from combination therapy targeting AR and AKT.

## Introduction

The *TMPRSS2:ERG* fusion (*T:E* fusion) puts *ERG* under the androgen receptor (AR) regulated expression of *TMPRSS2*, and is an early truncal alteration found in about half of prostate adenocarcinoma (PCa) cases in men of European ancestry. It can occur through an interstitial deletion of genes located between *TMPRSS2* and *ERG* on chromosome 21, or through a genomic rearrangement that preserves the intervening genes. Some early studies had indicated that tumors with the *T:E* fusion (or possibly with *T:E* fusion associated with an interstitial deletion) were more aggressive, but further studies have not found any clear differences in prognosis or responses to therapy (1, 2). The importance of ERG expression in driving PCa is supported by studies in the *T:E* fusion positive VCaP cell line, where RNAi mediated downregulation of ERG impairs cell growth and invasion (3, 4). Moreover, transgenic overexpression of ERG in mouse prostate causes increased proliferation, and in combination with loss of one *PTEN* allele results in prostatic intraepithelial neoplasia (PIN) or invasive PCa in aged mice (5–8). In varying contexts ERG in PCa has been found to activate proteins/pathways including EZH2, EMT (through ZEB1, ZEB2, and ILK), Wnt signaling, NFkB, SOX9, and YAP1 (3, 4, 8–16). More recently, it has become clear that ERG expression has marked global effects on gene expression in PCa cells, and in particular on the AR cistrome and transcriptome, where it directly interacts with AR and functions to maintain or expand AR signaling and luminal epithelial lineage (16–23).

The requirement for *PTEN* loss to drive PCa progression in mice overexpressing ERG is consistent with an association between *T:E* fusion and *PTEN* loss in human PCa, with *T:E* fusion presumed to be an initiating event and subsequent *PTEN* loss driving progression. One suggested basis for this association is that ERG is needed to maintain AR functions related to luminal differentiation in the context of *PTEN* loss (17–23). Indeed, one study found that ERG overexpression in PCa from *Pten* deficient mouse prostate mitigated responses to both AR targeted therapy and combination therapy with a phosphatidylinositol 3-kinase (PI3K) inhibitor (BEZ3235), and this was associated with maintenance of AR target gene expression (19). Conversely, another study found that ERG could suppress PI3K signaling and subsequent AKT activation, and identified ERG repression of insulin receptor substrate 2 (IRS2) expression as a possible mechanism (24). In this case, *PTEN* loss may be required as a secondary event to increase PI3K signaling and compensate for the repressive effects of ERG.

Inositol polyphosphate-4-phosphatase type II B (INPP4B) is another phosphatase that negatively regulates PI3K signaling and has been identified as a tumor suppressor in triple negative breast cancer (25, 26). Notably, analysis of TCGA data shows that *T:E* fusion in PCa is also strongly associated with decreased expression of *INPP4B* (24). In addition to negative regulation of PI3K signaling, INPP4B has further effects on signaling pathways through regulation of endosomal trafficking of proteins including receptor tyrosine kinases (RTKs) (27–29). However, the role of ERG in its regulation and the functional significance of its decreased expression in *T:E* fusion tumors remain to be determined.

Previous studies had shown that AR can suppress AKT activity by increasing expression of FKBP5, which is a scaffold for the AKT phosphatase PHLPP1 (30, 31). This provides one mechanism through which AR targeted therapies may increase PI3K/AKT signaling, although a recent study indicates that PHLPP1 and PHLPP2 do not functions as AKT phosphatases (32). AR can also positively regulate expression of *INPP4B*, so that AR inhibition may further increase PI3K signaling by decreasing INPP4B (33). These stimulatory effects on PI3K may partially mitigate responses to AR inhibition, and have provided the rationale for clinical trials combining inhibition of AR and PI3K/AKT signaling (34). Notably, as ERG in *T:E* fusion PCa is driven by AR, its expression is decreased in response to AR inhibition, which may further enhance PI3K signaling and mitigate therapeutic responses. In this study we further assessed ERG regulation of *IRS2*, *INPP4B*, and PI3K signaling, and tested the hypothesis that AR inhibition further enhances PI3K signaling in *T:E* fusion positive PCa.

## Results

### *T:E* fusion is associated with decreased *INPP4B* expression independently of *PTEN* status

While *PTEN* loss is associated with *T:E* fusion, *PTEN* is not altered in approximately half of cases (**Figure S1A**) and its expression is not reduced in these cases where the gene is intact (**Figure 1A, S1B**). In contrast, although the *INPP4B* gene is rarely altered in primary PCa (**Figure S1A**), its mRNA levels are globally reduced in *T:E* fusion positive cases (**Figure 1B**). Consistent with this reduction, reversed phase protein analysis of PCa tumors in TCGA shows that INPP4B is the most reduced protein in *T:E* fusion positive versus negative PCa (**Figure S1C**).

**Figure 1.**
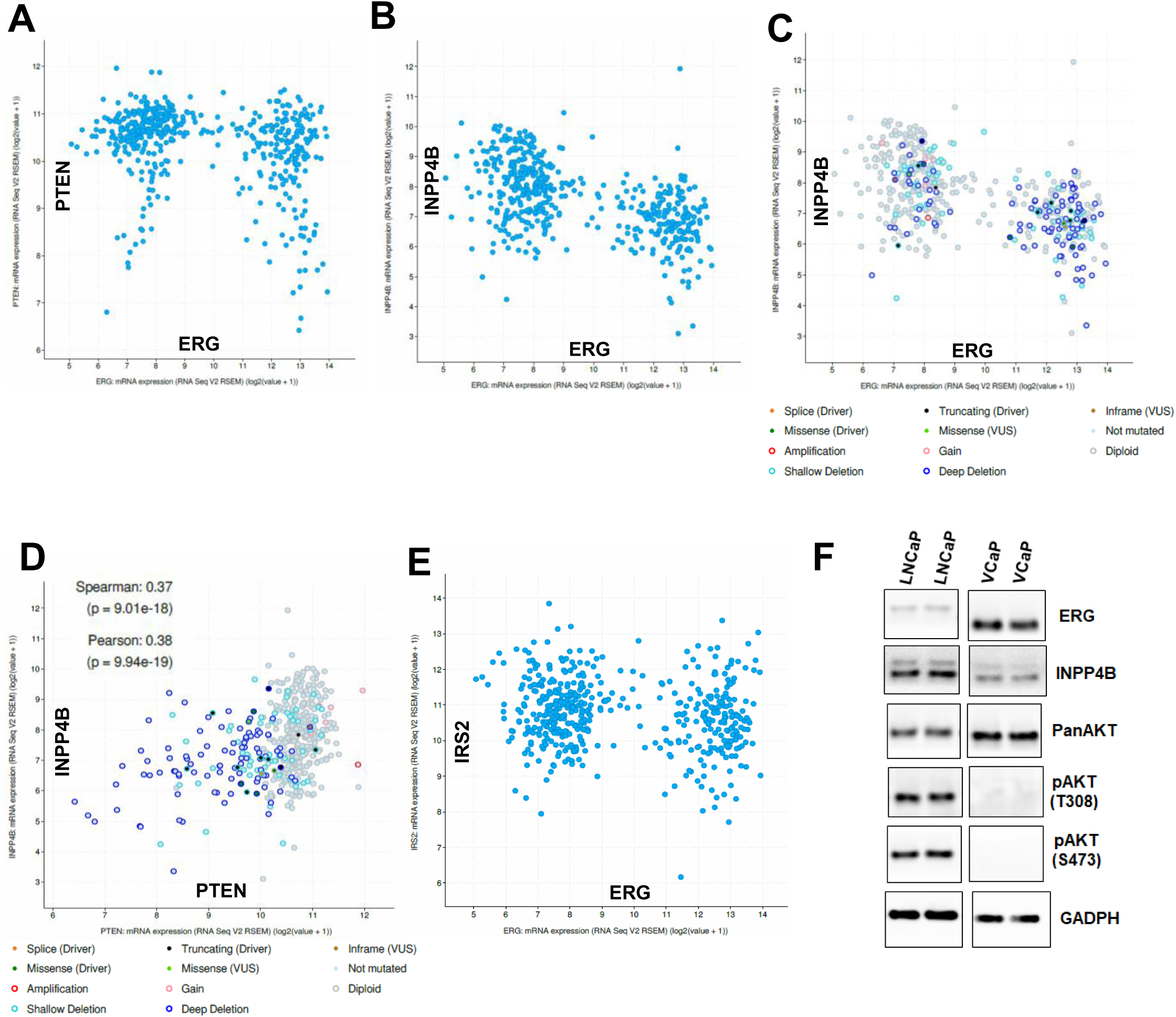
INPP4B mRNA is negatively correlated with *T:E* fusion independently of *PTEN* status. **(A)** Correlation between ERG and PTEN mRNA in TCGA primary PCa. **(B)** Correlation between ERG and INPP4B mRNA in TCGA primary PCa. **(C)** Correlation between ERG and INPP4B mRNA, and *PTEN* genomic alterations, in TCGA primary PCa. **(D)** Correlation between PTEN and INPP4B mRNA, and PTEN genomic alterations, in TCGA primary PCa. **(E)** Correlation between ERG and IRS2 mRNA in TCGA primary PCa. All analyses were done on cBioPortal. **(F)** Duplicate cultures of LNCaP and VCaP cells were lysed and immunoblotted as indicated.

INPP4B dephosphorylates phosphatidylinositol 3,4-bisphosphate (PI(3,4)P_2_), which similarly to PI(3,4,5)P_3_ mediates the binding to and activation of AKT and additional proteins with pleckstrin homology domains. Therefore, similarly to *PTEN* loss, the downregulation or loss of *INPP4B* can enhance AKT activation. However, they are not redundant. While PTEN primarily inhibits AKT activation at the plasma membrane, INPP4B targets endosomal AKT activation and also regulates the endosomal trafficking of multiple proteins including receptor tyrosine kinases (27–29). Notably, consistent with nonredundant functions, this reduced *INPP4B* expression is independent of *PTEN* status, as it is similarly reduced in the *T:E* fusion-positive tumors whether *PTEN* is lost or intact (**Figure 1C**).

Also consistent with nonredundant functions, there is a positive correlation between *PTEN* and *INPP4B* expression that is independent of *T:E* fusion status (**Figure 1D**). As shown previously (24), *IRS2* expression is lower in *T:E* fusion positive tumors (**Figure 1E**), although this decrease is not as marked as the decrease in *INPP4B*. We also examined INPP4B in the VCaP PCa cell line, which is *T:E* fusion positive and *PTEN* intact. Compared to LNCaP PCa cells (*T:E* fusion negative, *PTEN* deficient), VCaP has markedly lower INPP4B (**Figure 1F**). As expected, AKT activation, as assessed by phosphorylation at T308 and S473, is greater in the LNCaP cells.

### ERG suppresses PI3K signaling in PCa cells in vitro

To directly assess effects of ERG on PI3K signaling we used RNAi to decrease expression of ERG in *T:E* fusion-positive VCaP PCa cells. Consistent with a previous study (24), shRNA-mediated suppression of ERG increased PI3K signaling based on increased phosphorylation of AKT and S6 (**Figure 2A**). There was also a small increase in ERK phosphorylation. Acute suppression of ERG with siRNA similarly increased AKT and S6 phosphorylation (**Figure 2B**).

**Figure 2.**
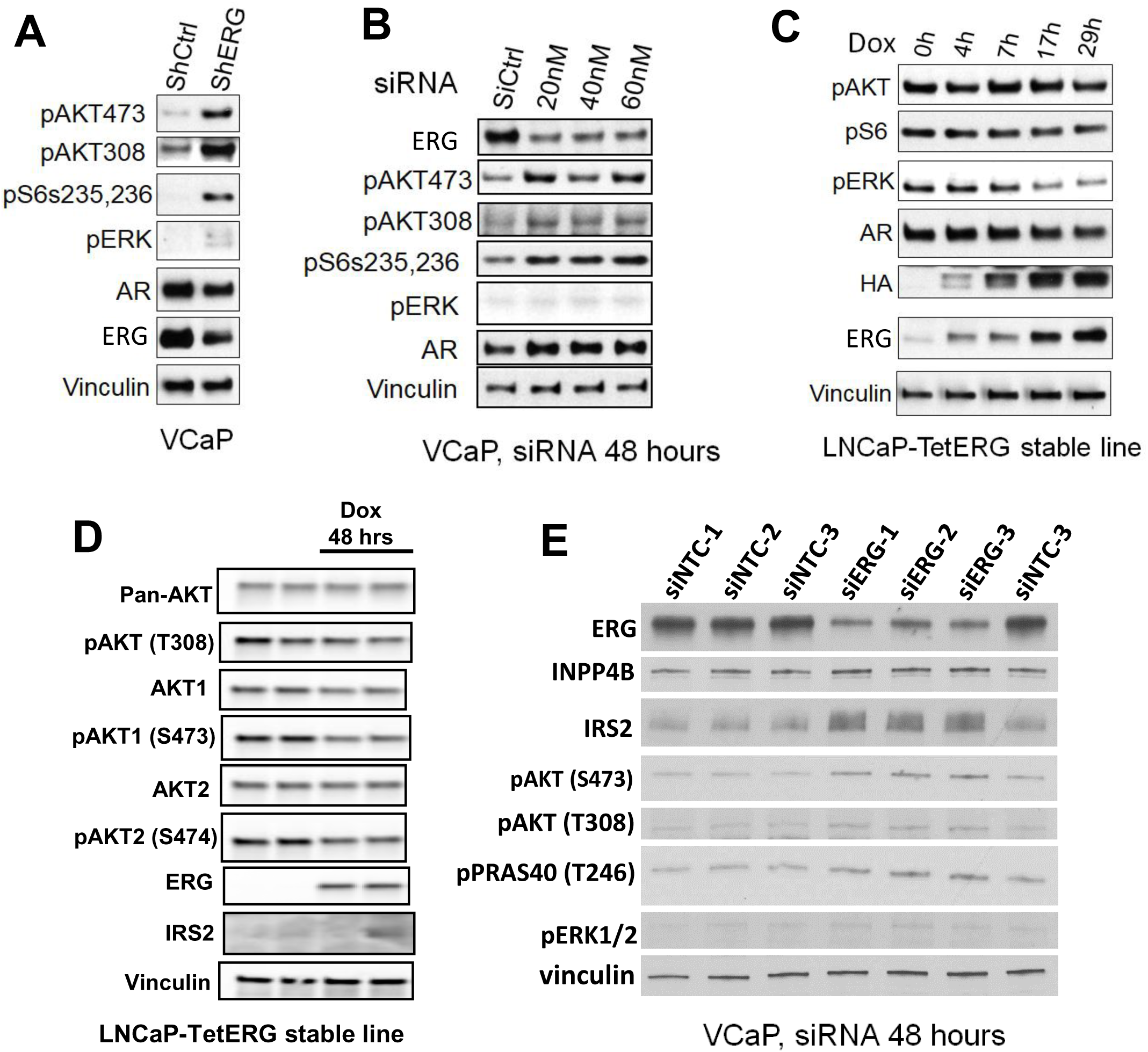
ERG suppresses AKT activity and IRS2 expression in PCA cells. **(A)** VCaP cells stably expressing an ERG shRNA versus a nontargeting control shRNA were assessed by immunoblotting as indicated. **(B)** VCaP cells were treated for 2 days with siRNA targeting ERG or a nontargeted control and assessed by immunoblotting as indicated. **(C)** LNCaP cells stably expressing a DOX-inducible HA-tagged ERG were treated with DOX over a time course and assessed by immunoblotting as indicated. **(D)** LNCaP cells stably expressing a DOX-inducible HA-tagged ERG were treated with DOX for 48 hours and assessed by immunoblotting. **(E)** VCaP cells were treated for 2 days with siRNA targeting ERG or nontargeted control and assessed by immunoblotting as indicated.

Conversely, we assessed the effects of inducing ERG expression in *T:E* fusion-negative LNCaP PCa cells. For this we generated a tet-operator (tetO) regulated vector (pTET-Splice) expressing N-terminal HA-tagged ERG, using human ERG with deletion of amino acids 1-39 to mimic the product generated from the most common *T:E* fusion (pTET-ERG) (**Figure S2A, B**). This was stably transfected into LNCaP cells expressing pcDNA6/TR for doxycycline (DOX)-regulated expression. The LNCaP cells are *PTEN* deficient and hence have high basal PI3K/AKT signaling (**Figure 2C**). DOX-mediated induction of ERG for up to 29 hours caused a small decrease in AKT and S6 phosphorylation, and a clear decrease in ERK phosphorylation (**Figure 2C**). The modest decrease in phosphorylation of AKT was more apparent after 48 hours of DOX induction, and was observed in both the AKT1 and AKT2 isoforms (**Figure 2D**).

Notably, ERG induction in LNCaP cells did not decrease expression of IRS2 (**Figure 2D, S2C**), indicating that the modest acute effects of ERG on AKT and ERK are not mediated via IRS2. We similarly examined effects of acutely inducing ERG in the 22RV1 PCa cell line, which is *T:E* negative and *PTEN* intact. In two independent lines we also observed only small and inconsistent effects of ERG induction on AKT activation and IRS2 expression (**Figure S3**).

### ERG suppresses IRS2 directly and INPP4B indirectly in *T:E* fusion positive cells

We next examined effects of depleting ERG on IRS2 and INPP4B in *T:E* fusion positive VCaP cells. Consistent with a previous report (24), ERG knockdown increased the expression of IRS2 protein (**Figure 2D**). In contrast, there was not a clear increase in INPP4B protein. RNA-seq analysis confirmed a ∼2.5-fold rise in IRS2 mRNA upon ERG siRNA treatment, whereas INPP4B was not changed (**Table S1**). Analysis of available ERG ChIP-seq data in VCaP cells showed an ERG binding site at the promoter of *IRS2*, but not *INPP4B*, further indicating that ERG directly regulates IRS2 and not INPP4B (**Figure S4A, B**). AR ChIP-seq further shows that this ERG binding site overlaps a broad AR binding site present in VCaP cells, but not in *T:E* negative LNCaP cells (**Figure S4A**). This is consistent with previous results showing that ERG opens cryptic AR binding sites in many genes, so these come under joint ERG and AR regulation (16).

As noted above, the acute induction of ERG in LNCaP and 22Rv1 cells did not cause a clear decrease in IRS2, indicating that the ERG suppression of IRS2 in *T:E* positive tumors requires further adaptations. Interestingly, although ERG knockdown increases IRS2 mRNA in VCaP cells, it did not increase H3K27Ac at the *IRS2* promoter (**Figure S4A**), suggesting ERG is repressing through a mechanism distinct from modulation of histone acetylation.

To assess more broadly the effects of ERG knockdown we used gene set enrichment analysis (GSEA). PI3K-AKT-MTOR signaling was only modestly increased in ERG knockdown cells (**Figure S5A,B**), while the most enriched gene set after ERG knockdown was Androgen Response followed by Fatty Acid Metabolism (which may be androgen-regulated) (**Figure S5A, C, D**). This result is somewhat surprising as ERG can maintain or expand AR signaling in the setting of *PTEN* loss, and may reflect that VCaP is *PTEN* intact. Conversely, Hallmark gene sets including MYC targets and E2F targets were markedly decreased in response to ERG knockdown (**Figure S5A, D**), consistent with an oncogenic function of ERG in these cells. Together these results support the conclusion that ERG directly suppresses IRS2 expression in *T:E* fusion tumors. In contrast, these results also indicate that the decreased expression of *INPP4B* in *T:E* fusion tumors is not direct, and that it is an adaptation that is selected, at least in part, to compensate for suppressive effects of ERG on PI3K signaling, and potentially ERK signaling.

### AKT activation is inversely associated with ERG expression in mouse model

We next used a mouse model to assess ERG function in an in vivo context. We used the insert from the pTET- ERG vector to generate transgenic mice expressing HA-tagged human ERG (amino acids 1-39 deleted) under the control of the tetracycline operator. We had previously developed transgenic mice with a probasin-driven reverse tetracycline transactivator (rtTA, tet-on) that can drive prostate specific expression of tetracycline operator-driven transgenes (35). These mice were crossed with the pTET-ERG transgenic mice to obtain DOX-stimulated expression of ERG in the prostate. We initially sacrificed a small cohort of mice at ∼4-5 months after DOX induction, and isolated protein from each prostate lobe. By immunoblotting we found human ERG expression in the ventral lobe, which we confirmed by qRT-PCR (**Figure S6A**). By IHC we further confirmed DOX-driven human ERG expression in ventral prostate (**Figure S6B**), and in two pTET-ERG lines confirmed that expression was dependent on the rtTA (**Figure S6C**). Consistent with previous studies showing a modest effect of ERG overexpression in mouse prostate (5, 6), induction of ERG for as long as 16 months caused only mild hyperplasia (**Figure S6D**).

Next, we crossed these mice onto a *Pten* haploinsufficient background (*Pten^-/+^*) (5). In these mice we then induced ERG expression by feeding mice with both DOX food and water to further increase ERG expression (**Figure S7A, B**). We identified areas of PCa after ∼11 months of DOX induction (**Figure 3A**). Notably, ERG expression was heterogeneous, with much lower ERG expression in the areas with invasive tumor (**Figure 3B**).

**Figure 3.**
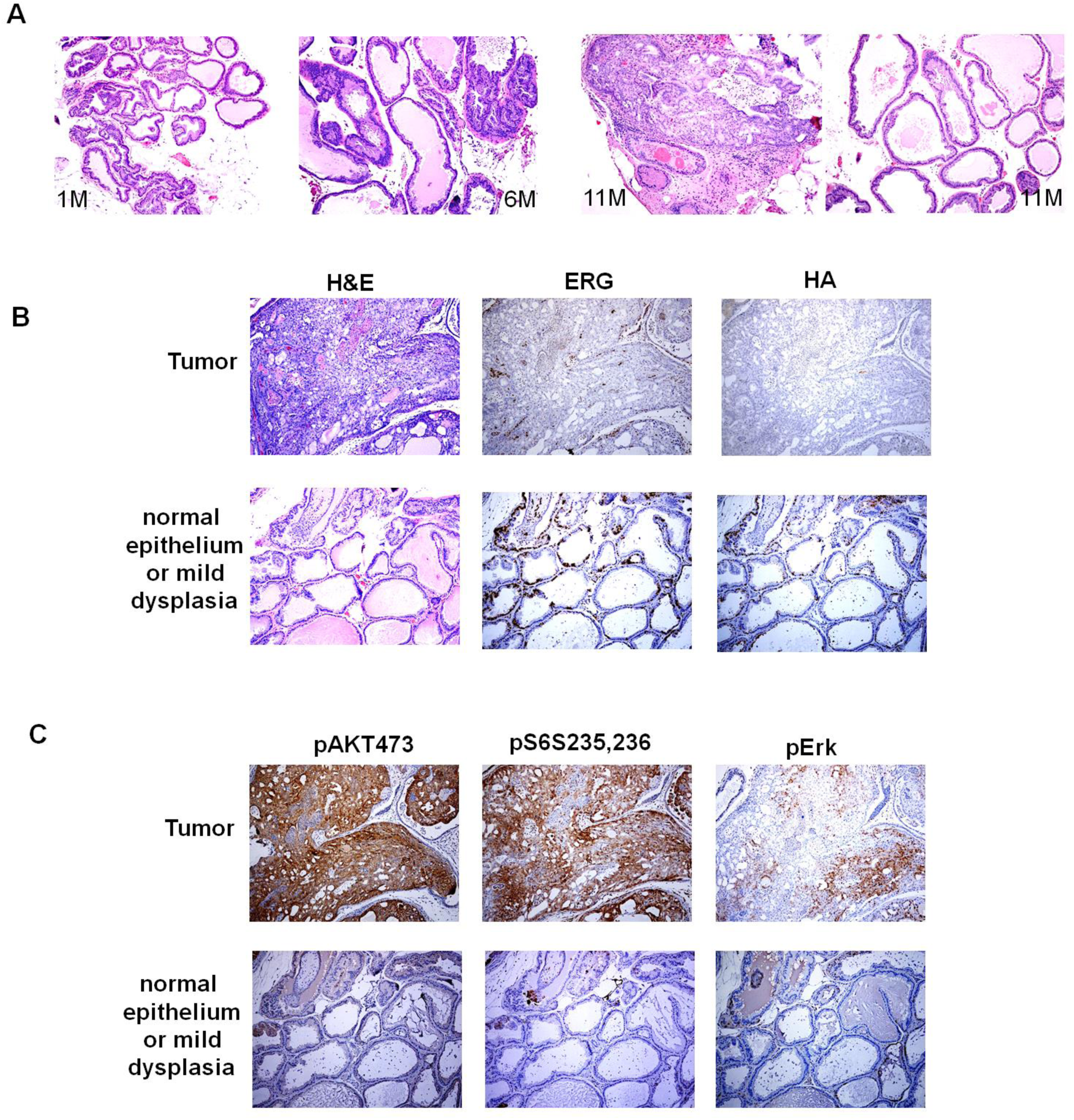
PCa development in probasin-rtTA; tetO-ERG; *Pten^-/+^* mice. **(A)** Histology in mice treated for 1 – 11 months with DOX, showing foci of PCa in mouse treated with DOX for 11 months. **(B)** FFPE sections from representative areas with or without tumor were stained for ERG or the HA tag on ERG. Lower ERG expression is seen in areas with invasive tumor versus areas showing only dysplasia. **(C)** Higher pAKT and pS6 in areas with invasive tumor versus areas showing only dysplasia.

Moreover, in those tumor areas with low ERG, we found elevated expression of pAKT(473), pS6(S235,236), and to a lesser extent pERK, reflecting activation of the PI3K/AKT pathway (**Figure 3C**). These observations were consistent with ERG suppression of PI3K/AKT signaling. The decrease in ERG in these areas may be due in part to a decrease in AR, which is driving the rtTA in these mice (**Figure S8A**). Expression of another AR-driven gene, *Tmprss2*, was also decreased in these tumor areas with low ERG (**Figure S8B**). These findings indicate that while ERG is oncogenic, higher levels may be tumor suppressive, possibly due to suppression of PI3K/AKT signaling.

### ERG-regulated genes in probasin-rtTA; tetO-ERG; *Pten^-/+^* mice

To identify ERG-regulated genes that may be contributing to PCa development in this model, we carried out time course experiments with both short-term ERG induction and long term induction plus short term DOX withdrawal in the probasin-rtTA; tetO-ERG; *Pten^-/+^* (*ERG;Pten^+/-^*) mice. For the short-term induction, mice were treated with DOX or vehicle for 3 or 6 days. Ventral prostate epithelium was then laser capture microdissected and transcriptome profiles were obtained using Affymetrix microarrays. The level of ERG induction based on the microarrays was similar at 3 and 6 days (∼25% increase), so expression data from both days were pooled. There were 5 Hallmark gene sets that were significantly (FDR <0.25) increased and 16 decreased (**Figure S9A**). The MTORC1 Signaling and PI3K AKT MTORC Signaling were amongst the highly suppressed gene sets (NES -3.34 and -2.64 respectively) (**Figure S9B, C**). Notably, this suppression could not be attributed to IRS2, as it was not decreased by ERG induction (ratio induced/uninduced ∼1.18). INPP4B was increased (ratio induced/uninduced ∼1.16), indicating it may acutely contribute to decreased PI3K signaling in these mice.

Interestingly, the most altered gene involved in phosphatidylinositol metabolism was PIKFYVE (phosphoinositide kinase) (ratio induced/uninduced 0.71). This kinase phosphorylates the D-5 position in phosphatidylinositol and phosphatidylinositol-3-phosphate on endosomes, the latter generated by INPP4B, to promote lysosomal targeting. The second most altered gene in phosphatidylinositol metabolism was phosphoinositide-3-kinase regulatory subunit 3 (PIK3R3, ratio induced/uninduced 0.74), which may contribute directly to a decrease in PI3K activity in these mice. Analysis of TCGA PCa data show that PIK3R3 is significantly decreased in *T:E* fusion tumors (**Figure S9D**), indicating this may also contribute to ERG-mediated suppression of PI3K signaling in patients.

We next used RNA-seq to assess the effects of discontinuing DOX in mice with established tumors. There were no Hallmark gene sets altered at FDR <0.25. Nonetheless, IRS2 was ∼4-fold higher in tumors after DOX discontinuation, consistent with it being suppressed by ERG in the established tumors. Levels of INPP4B prior to and after DOX discontinuation were too low for reliable assessment. Interestingly, the lncRNA H19 was the most increased transcript in the DOX induced tumors versus levels after DOX withdrawal (**Figure 4A**). H19 was also increased by short-term DOX induction (although only about 2-fold versus the >100-fold increase in the long-term DOX induced tumor samples) (not shown), indicating that ERG is inducing H19. Notably, *H19* and *IGF2* are co-regulated in human and mouse by genomic imprinting through methylation at the locus between *H19* and *IGF2*, and we also observed increased IGF2 in the DOX-induced tumors versus the samples after DOX withdrawal (**Figure 4A**). Marked ERG-mediated increases in IGF2 and H19 mRNA were also observed in a previous independent study using stable probasin-ERG transgenic mice, but the significance of this increase with respect to decreased PI3K/AKT signaling is not clear (13) (**Figure 4B**).

**Figure 4.**
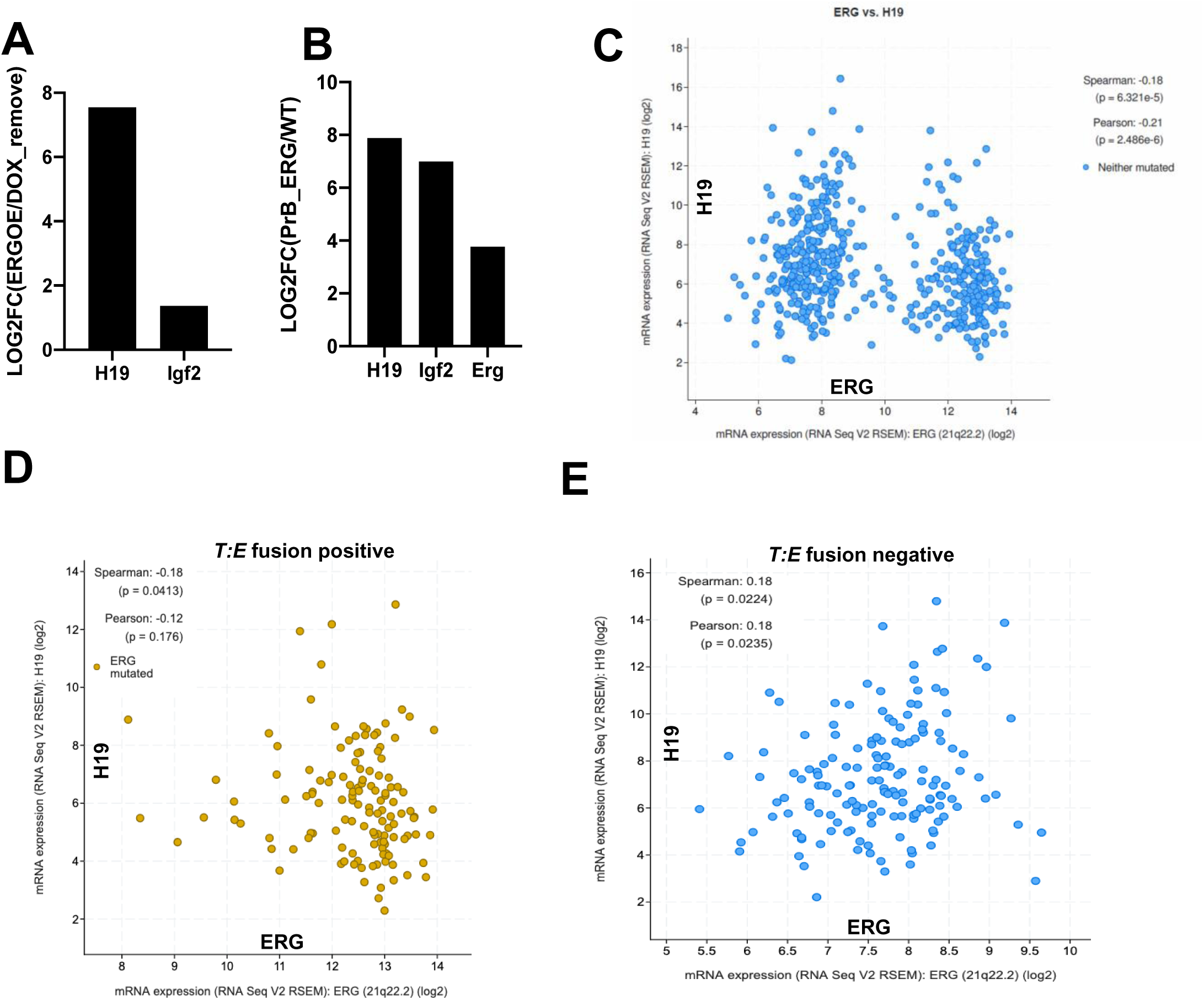
H19 lncRNA regulation by ERG in mouse model and human PCa. **(A)** H19 and IGF2 expression in mice with long-term DOX induction versus after DOX withdrawal. **(B)** H19, IGF2, and ERG expression in mouse prostate expressing probasin-ERG transgenes versus wild-type (from reference 13). **(C)** Correlation between ERG and H19 mRNA in TCGA primary PCa. **(D)** Correlation between ERG and H19 mRNA in *T:E* fusion positive tumors in TCGA primary PCa. **(E)** Correlation between ERG and H19 mRNA in *T:E* fusion negative tumors in TCGA primary PCa. Correlations were assessed in cBioPortal.

While H19 appears to be a major target of ERG in mouse, analysis of gene expression data in human primary PCa shows that H19 is instead decreased in *T:E* fusion positive tumors (**Figure 4C**). Interestingly, analyzing *T:E* fusion-positive and *T:E* fusion-negative tumors separately we find that H19 expression is positively correlated with ERG in *T:E* negative (but not positive) tumors, but the significance of this association is not clear (**Figure 4D, E**). Together these findings indicate that H19 is a major transcriptional target of ERG in this mouse PCa model, but not in human PCa. Overall these observations support the conclusion that ERG is repressing PI3K/AKT signaling in this mouse PCa model and in human PCa, although some distinct mechanisms may be involved.

### Castration enhances AKT activation in *T:E* fusion positive PDX model

As expression of ERG in *T:E* fusion positive tumors is driven by AR, it is markedly decreased in response to AR targeted therapies. Therefore, we next assessed the extent to which AR inhibition increases PI3K/AKT signaling in VCaP cells. As expected, treatment with increasing concentrations of an AR antagonist (enzalutamide) caused a progressive decrease in ERG and PSA, and further decreased INPP4B, which is AR regulated (**Figure S10**). This was associated with an increase in phosphorylation of AKT2, but no clear change in AKT1. Interestingly, there was also a decrease in PHLPP1 and PHLPP2 that may dephosphorylate AKT (30, 31), although a recent study indicates these do not function as AKT phosphatases (32). Finally, PTEN expression was unchanged.

To assess effects of AR inhibition in vivo, we examined a *T:E* fusion positive/*PTEN* negative PDX (BIDPC4) we derived from an omental metastasis in a patient with CRPC. PDXs were established subcutaneously in male *scid* mice and analyzed prior to castration and at ∼2 weeks post-castration. Castration decreased ERG (as expected) and caused a marked increase in phosphorylation of AKT and S6 (**Figure 5A**). To explore the role of AKT activity in this model, we generated short-term ex vivo cultures from BIDPC4 PDX cells. These cultures were treated with an AKT inhibitor (ipatasertib) in either medium containing fetal bovine serum (FBS) or androgen-depleted charcoal-stripped serum (CSS). Notably, cells in the CSS medium showed heightened sensitivity to AKT inhibition (**Figure 5B**). A similar effect was observed with a second AKT inhibitor, MK2206 (**Figure 5C**).

**Figure 5.**
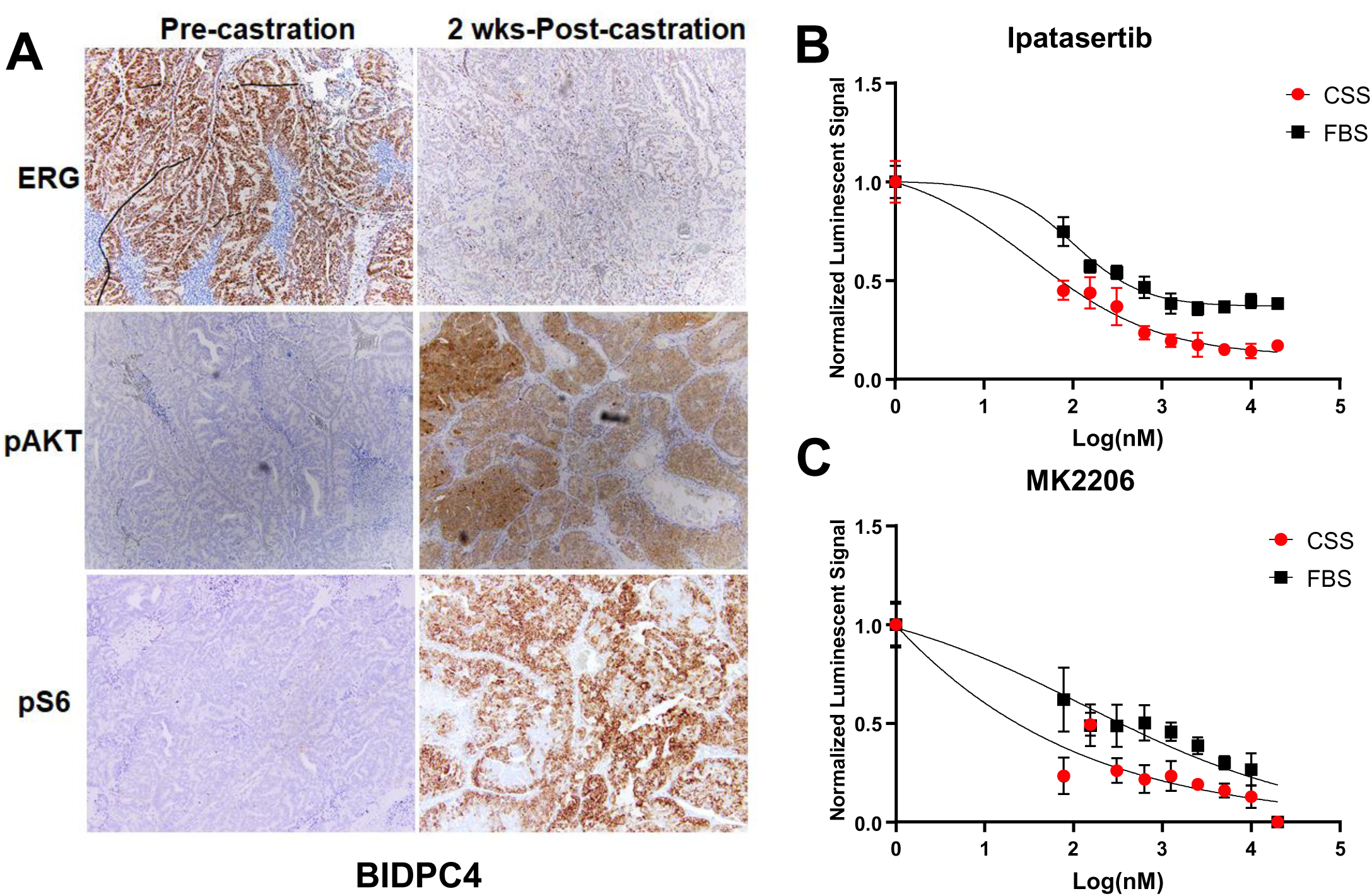
Androgen deprivation increases AKT activity and dependence in *T:E* fusion positive PDX. **(A)** *T:E* fusion positive BIDPC4 PDXs were established in male *scid* mice as subcutaneous tumors. Mice were sacrificed prior to or at ∼2 weeks post-castration. FFPE sections were stained for ERG, pAKT(473), and pS6 as indicated, and representative slides are shown. **(B)** Cells from BIDPC4 PDX were cultured in medium with 10% FBS (FBS) or in medium with 10% charcoal stripped FBS (CSS). Ipatasertib was then added at the indicated concentrations and cell recovery was assessed by Cell Titer Glo assay after 7 days. Luminescent readouts were normalized to the average luminescent signal of DMSO under the respective conditions, with 6 replicates per dose of drug. The graph illustrates the half-maximal inhibitory concentration (IC50) in the presence of FBS compared to CCS. Nonlinear regression curves were generated by a variable slope model. (**C**) Responses to MK2206 were assayed as in (B).

### AR signaling inhibition enhances AKT activation in *T:E* fusion positive clinical samples

We next examined available clinical data to assess the extent to which AR targeted therapies increase AKT activation in *T:E* fusion positive versus negative PCa. For this analysis we used genomic data and ERG mRNA levels to identify *T:E* fusion positive tumors in the TCGA data set of primary PCa (36) and in two CRPC data sets (37, 38) (**Figure S11A**). We then carried out single sample GSEA (ssGSEA) for each tumor using the Hallmark PI3K_AKT_MTOR gene set to determine whether AKT signaling was increased in the *T:E* positive CRPC cases relative to the TCGA primary tumors, but did not find a significant difference in the *T:E* positive or negative tumors (**Figure S11B**). We similarly examined a small series of cases that had matched RNA-seq data from CRPC tumors before and after treatment with AR inhibitors (39). While there was enrichment for the PI3K_AKT_MTOR gene set in some *T:E* positive cases, the increase was not significant and was similar to that in the *T:E* fusion negative cases (**Figure S11C**).

Notably, there is substantial restoration of AR activity in most CRPC cases, and we reported previously that this includes increased ERG expression *T:E* fusion positive CRPC (40). Indeed, in the matched untreated versus treated CRPC samples, ERG remained markedly elevated in the *T:E* fusion positive versus negative tumors (**Figure S11D**). Therefore, to further assess effects of AR targeted therapy on tumors prior to emergence of CRPC, we examined samples from a clinical trial of neoadjuvant intensive AR targeted therapy (NCT02430480) (41).

In this trial, patients with high-risk primary PCa were treated for 24 weeks with leuprolide in combination with enzalutamide, followed by radical prostatectomy (RP). Significantly, IHC for ERG showed that ERG was markedly decreased in the residual tumor in the RP specimens versus the pretreatment biopsies (**Figure 6A, B**). ERG mRNA was similarly greatly decreased in the post treatment *T:E* fusion positive tumors (**Figure 6B**). Analysis of the RNA-seq data by ssGSEA then showed that the PI3K_AKT_MTOR gene set was significantly increased (P = 0.020) in the post treatment versus matched pretreatment tumors (**Figure 6C**). This gene set was also modestly increased in some *T:E* fusion negative cases, but this was not significant (**Figure 6C**).

**Figure 6.**
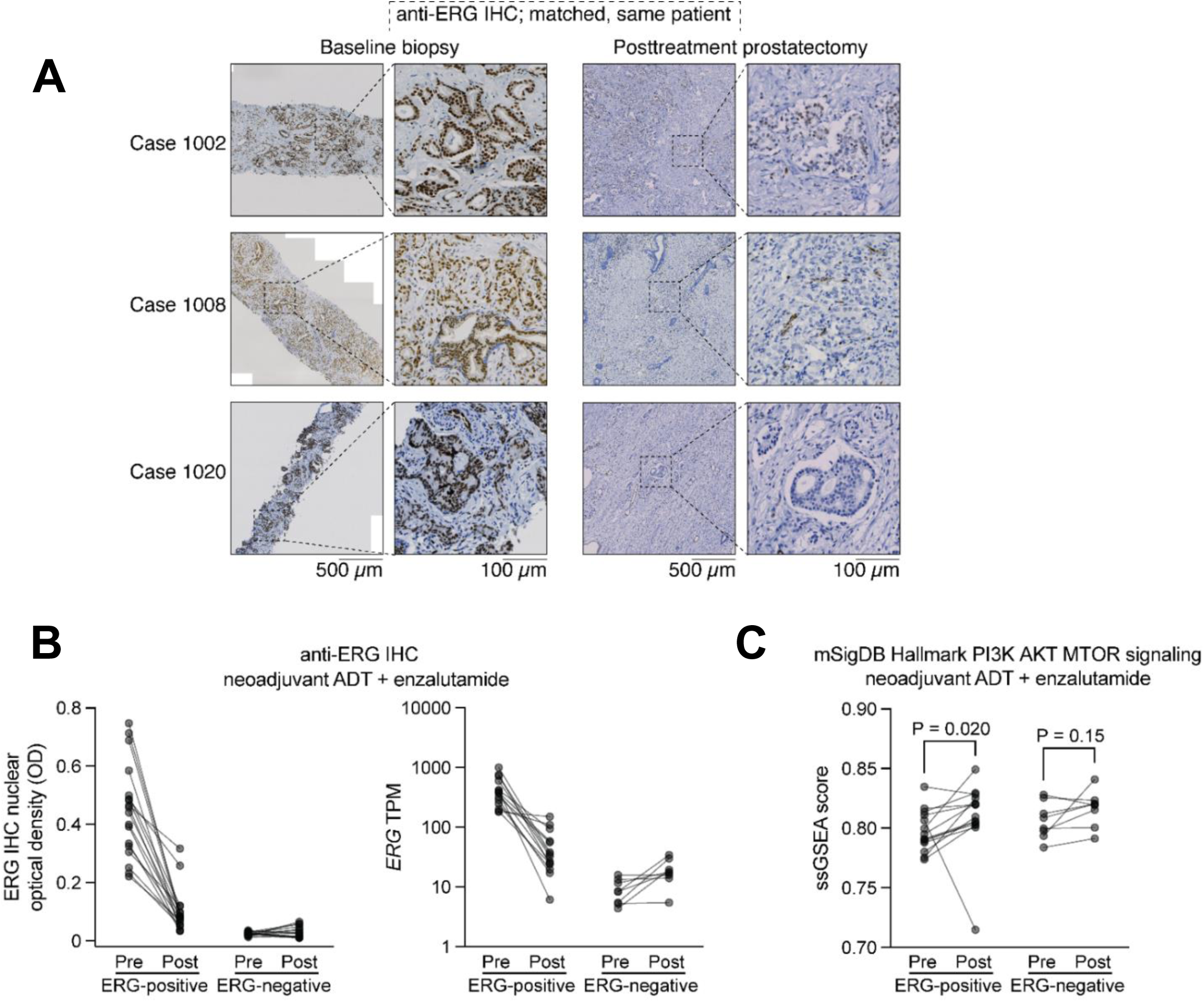
PI3K_AKT_MTOR activity in primary PCa before and after neoadjuvant AR-targeted therapy. **(A)** FFPE sections of tumor from matched pairs of baseline biopsy and posttreatment radical prostatectomy specimens stained with anti-ERG antibodies, from three representative *T:E* fusion-positive cases. Bar: 500 µm; inset bar: 100 µm. **(B)** ERG levels in matched pairs as determined by (left) anti-ERG IHC quantified by machine-guided image analysis or (right) RNA-seq. **(C)** Single-sample GSEA for the Hallmark PI3K_AKT_MTOR gene set in *T:E-*positive (left, n=14) and *T:E*-negative (right, n=8) tumors. Statistical significance determined using the Wilcoxon matched-pairs signed rank test.

As expected, *PTEN* loss was greater in the *T:E* fusion positive (13/16) versus negative (9/21) tumors (**Figure S11D**). This high frequency of *PTEN* loss in both groups may reflect selection for patients with high-risk tumors. ERG expression by IHC was markedly decreased by the neoadjuvant therapy in both the PTEN intact and deficient *T:E* fusion tumors (**Figure S11E**). Due to limited residual tumor in the *PTEN* intact *T:E* tumors we were not able to carry out RNA-seq to assess effects of the treatment on PI3K/AKT signaling. However, effects in the *T:E* negative tumors did not appear related to *PTEN* status (**Figure S11F**).

We also examined samples from a clinical trial of neoadjuvant leuprolide, abiraterone, and apalutamide administered for 24 weeks prior to RP in men with high-risk primary PCa (NCT02903368) (42). ERG IHC on diagnostic biopsies was used to identify cases that were *T:E* fusion positive versus negative. Diagnostic core biopsies and corresponding RP specimens containing residual tumor in the *T:E* positive and *T:E* negative cases were then stained for phospho-AKT (**Figure 7A**). Notably, phospho-AKT staining was variable in the biopsies. Therefore, to take into consideration factors other than ERG that may influence PI3K signaling, we quantified the difference in phospho-AKT staining between the biopsies and RP specimens for each case. Staining in the RP specimens was greater than in the matched biopsies in over half of the *T:E* positive cases versus only 2 of the *T:E* negative cases (**Figure 7B**), and this increase in the *T:E* fusion positive versus negative residual tumors was significant at P < 0.01 (**Figure 7C**). Taken together these findings indicate that AR targeted therapies may increase PI3K/AKT pathway activity by several mechanisms, with decreased ERG expression in *T:E* fusion tumors being a major mechanism in this tumor subset (**Figure S12**).

**Figure 7.**
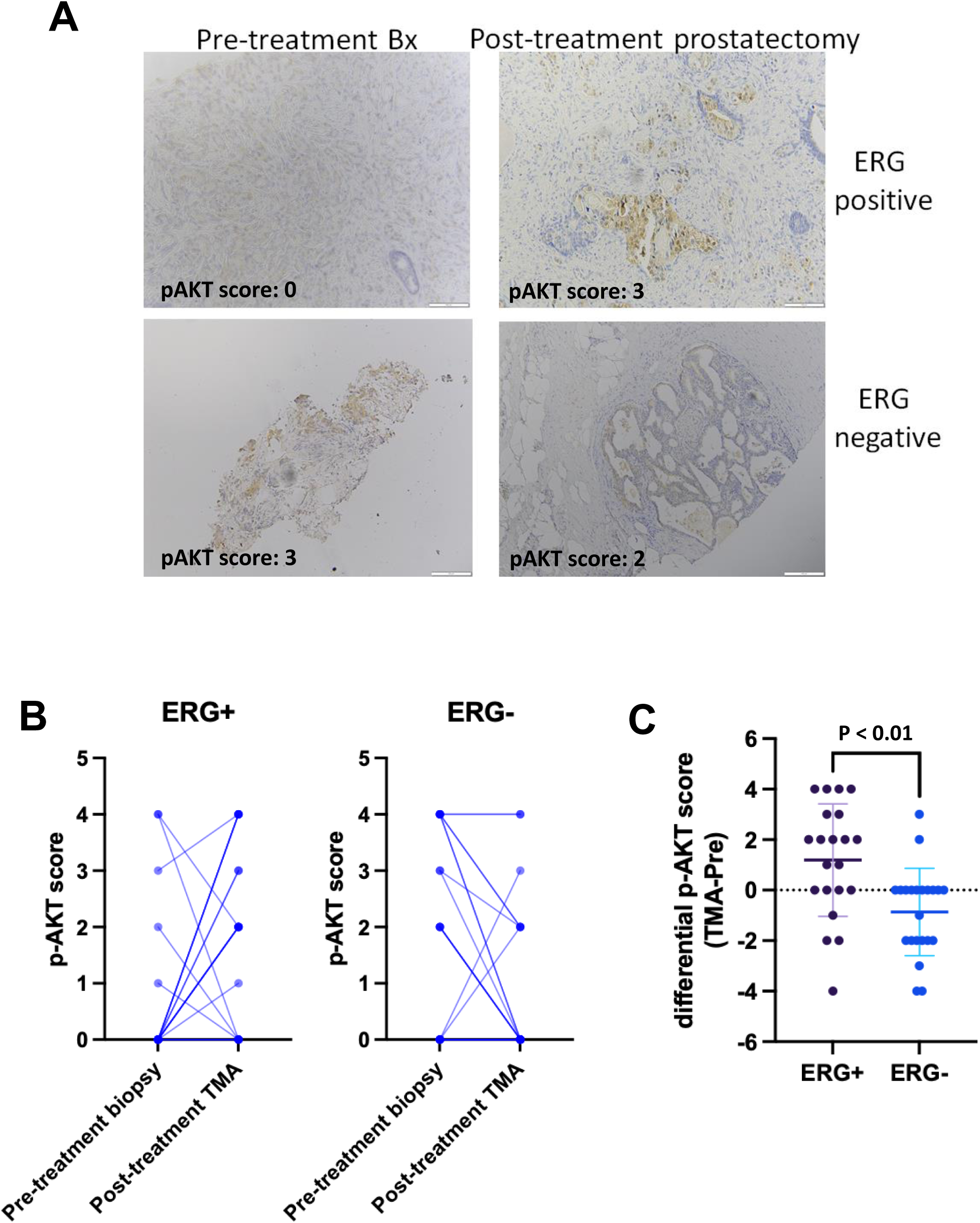
AKT phosphorylation prior to and post-neoadjuvant AR targeted therapy. **(A)** FFPE sections of tumor in pretreatment biopsies and in radical prostatectomy specimens from a *T:E* fusion positive and negative case were stained for pAKT(473). Samples were scored based on fraction of cells staining. Pretreatment biopsy scores for the *T:E* fusion positive and negative tumors shown here were 0 and 3, respectively. Scores for the corresponding post-treatment tumors were 3 (*T:E* fusion positive) and 2 (*T:E* fusion negative). **(B)** pAKT scores for each pre-treatment and corresponding post-treatment tumor in *T:E* fusion positive (left panel) and negative (right panel) cases. Note that darker lines reflect multiple cases with the same pre- and post-treatment scores. **(C)** For each patient the pAKT score in the pretreatment biopsy was subtracted from the score in the corresponding post-treatment tumor to give a differential pAKT score. ** p < .05.

## Discussion

*T:E* fusion is frequently associated with *PTEN* loss and is strongly associated with decreased *INPP4B* expression. The basis for these associations remains to be established, but one possibility is that ERG may suppress PI3K/AKT signaling and that decreased expression of PTEN and INPP4B may compensate for this suppression. Our finding that ERG knockdown enhances PI3K signaling in *T:E* fusion positive VCaP cells supports this conclusion and is consistent with results in a previous study (24). Also consistent with this previous study, we found that ERG knockdown in VCaP cells increased IRS2, supporting ERG repression of IRS2 as a mechanism contributing to ERG suppression of PI3K signaling. In contrast, despite the strong negative correlation between ERG and INPP4B levels in clinical samples, ERG knockdown did not increase INPP4B mRNA or protein, suggesting the decreased expression of INPP4B is indirect and reflects potent selective pressure to decrease INPP4B function. Results in mice with DOX-regulated induction of ERG further support the conclusion that ERG suppresses PI3K signaling, although mechanisms driving this decrease and subsequent adaptations in mouse may in part be distinct. Interestingly, while ERG appears to be directly suppressing IRS2 in *T:E* positive tumors, IRS2 is not suppressed by the acute induction of ERG, indicating that the ERG suppression of IRS2 is dependent on further adaptations.

Significantly, INPP4B expression is strongly and broadly decreased in *T:E* fusion positive PCa, including in cases with genomic *PTEN* loss. This indicates that *PTEN* loss alone does not fully compensate for effects of ERG induction, and that decreasing INPP4B may have distinct effects that are required to support the growth of *T:E* fusion tumors. Indeed, PTEN suppresses AKT activation primarily by decreasing P(3,4,5)P_3_ (converting it to (P4,5)P_2_) at the plasma membrane, although it may in some settings also target P(3,4)P_2_ (43). Its loss results in increased AKT recruitment to the plasma membrane and subsequent AKT phosphorylation and activation. *PTEN* loss is common in primary PCa, and its frequency is increased in metastatic CRPC, which is presumed to reflect increased AKT activation and subsequent increases in tumor cell survival and invasion.

In contrast, INPP4B-mediated dephosphorylation of PI(3,4)P2 primarily regulates AKT activation intracellularly, with one study indicating that INPP4B loss may selectively increase activation of the AKT2 isoform (28). Moreover, the PI(3)P generated by INPP4B on endosomal membranes can enhance lysosomal targeting of signaling proteins including EGFR and GSK3β to downregulate receptor tyrosine kinase signaling and to enhance Wnt signaling, respectively (27, 44, 45). To the extent that *T:E* fusion tumors need to compensate for decreased IRS2, this former increase in receptor tyrosine kinase signaling may be an important factor driving selection for decreased INPP4B. Further studies are needed to address the extent to which loss of these or other INPP4B functions drive selection for its downregulation in *T:E* fusion positive PCa, and whether this results in novel therapeutic vulnerabilities.

Previous studies have shown that AR-targeted therapies can increase PI3K signaling through mechanisms including downregulation of FKBP5 and of INPP4B (30, 31, 33). As ERG is AR regulated in *T:E* fusion positive tumors, we hypothesized that AR inhibition, and subsequent decreased expression of ERG, would in particular enhance AKT activation in these tumors. Indeed, we found that AKT activity and dependence was increased after castration in a *T:E* fusion positive PDX model. We then used ssGSEA to assess for increases in PI3K/AKT signaling in *T:E* positive versus negative CRPC, but did not find a significant increase. However, while ERG expression is decreased acutely after AR targeted therapies, it is substantially restored in CRPC. Therefore, we next examined matched samples from two neoadjuvant trials of intensive AR targeted therapy. We confirmed that ERG was markedly decreased in the post treatment residual tumor, and found that PI3K/AKT activity was significantly increased in the *T:E* fusion positive versus negative post treatment cases. Based on these findings, we suggest that increased AKT activity due to decreased ERG mitigates the efficacy of AR-targeted therapies to a greater extent in *T:E* fusion PCa, and that men with *T:E* fusion-positive tumors may most benefit from the addition of an AKT inhibitor to AR targeted therapy.

Notably, results of a phase 3 trial of abiraterone combined with an AKT inhibitor (ipatasertib) in CRPC (IPATential150 trial) indicated that there was radiographic progression free survival (rPFS) benefit in patients with *PTEN*-deficient tumors (34). As *PTEN* loss is associated with *T:E* fusion, it is possible that some of this benefit may reflect *T:E* fusion. A subsequent analysis of the IPATential150 trial (published while this manuscript was under revision) found that while there was an increase in median overall survival (OS) in men with *PTEN*-deficient tumors (from 29.8 months in the placebo control to 36.8 months in the ipatasertib arm), this did not reach statistical significance (46). Significantly, in an exploratory analysis this study found that median OS in *T:E* fusion tumors was improved from 30.8 months to 42.9 months by addition of ipatasertib, with no clear benefit in the *T:E* negative tumors.

These clinical trial findings further support the hypothesis that *T:E* fusion tumors may have increased dependence on AKT activation subsequent to AR targeted therapies. Moreover, we suggest that this benefit of adding an AKT inhibitor may be greater in patients with castration-sensitive PCa who are initiating AR targeted therapy. Notably, the phase 3 CAPItello-281 trial (NCT04493853) is assessing another AKT inhibitor (capivasertib) in combination with AR inhibition (GnRH agonist plus abiraterone and prednisone) in men with *PTEN* deficient metastatic castration-sensitive PCa. It will be of interest to determine whether the PTEN defiicinet tumors with *T:E* fusion have greater benefit in this trial. Finally, we suggest that *T:E* fusion tumors may have additional vulnerabilities, including those related to INPP4B downregulation, that may be exploited therapeutically.

## Methods

### Sex as a biological variable

Prostate cancer only occurs in males and is dependent on androgens. Therefore, the in vivo studies only used male mice.

### Cell lines and reagents

LNCaP and VCaP cells were obtained from ATCC and cultured in RPMI-1640/10% FBS or DMEM/10% FBS, respectively. Cells were used within 6 months of thawing, and were tested negative for Mycoplasma. siRNA targeting ERG and control SiCtrl were from Dharmacon (Thermo Fisher). shRNA plasmids targeting ERG and controls were from Santa Cruz. Transfections were performed using Lipofectamine 2000 (Invitrogen) according to manufacturer instructions. The pTET-ERG plasmid was made by inserting N-terminal HA-tagged ERG (with deletion of amino acids 1-39) into the tet-operator (tetO) regulated vector pTET-Splice (Invitrogen). LNCaP cells with DOX inducible ERG were then generated by transfecting the pTET-ERG vector into a LNCAP clone carrying pcDNA6/TR, followed by selection.

### RNA interference

ERG knockdown were performed in VCaP cells. Briefly, VCaP cells were transfected with 20, 40 and 60 nmol/L ON-TARGETplus siRNAs (Dharmacon) with Lipofectamine RNAiMAX (Thermo Fisher Scientific, 13778–075). At 48 hours after transfection, cells were harvested for protein purification and RNA purification.

### RNA isolation and qRT-PCR

Total RNA was isolated with the RNeasy Mini Plus Kit (Qiagen, #74134). RNA purification was directly performed from 2D cultured cells. For RNA isolation from tissues, homogenization were performed using a tissue ruptor(Qiagen, cat No. 9002755). qRT-PCR was performed using standard SYBR Green reagents from the StepOnePlus Real-Time PCR system (Thermo Fisher Scientific, #11780200) or TaqMan One-Step RT-PCR reagents (Thermo Fisher Scientific, #4444434). Target mRNA expression was quantified using the ΔΔCt method and normalized to GAPDH expression.

### Transgenic mice

The tetO regulated HA-ERG was excised from the pTET-ERG vector and used for pronuclear microinjection in FVB mice (BIDMC Transgenic Facility). Founder lines determined by genotyping were then bred with transgenic mice we generated previously expressing a probasin-driven reverse tetracycline transactivator (rtTA, tet-on), which we showed could drive prostate specific expression of tetracycline operator-driven transgenes (35). Transgene expression was induced by feeding a rodent diet containing doxycycline chow (0.625g/kg, Harlan Tekland, Indianapolis, IN) alone or with drinking water containing 1g/L DOX in 1% dextrose. *Pten* deficient mice on a C57BI/6J background were generously provided by Dr. Pier Paolo Pandolfi. Prostates were dissected and snap-frozen for RNA/protein extraction or formalin-fixed for paraffin embedding.

#### PDX tumors

The BIDPC4 PDX was generated from an omental metastasis in a patient with *T:E* fusion CRPC. Exome sequencing shows in addition the following: *AR* amplification, *PIK3CA* H419_L422del, *PTEN* loss, *MYC* amplification, *FAS* loss, *TP53* G245S. Male 5–6 weeks old ICR SCID mice were purchased from Taconic and housed in the Animal Research Facility at Beth Israel Deaconess Medical Center (BIDMC, Boston, MA). The PDX was initially established by inoculating human biopsies both subcutaneously and by renal graft with 50% Matrigel. Upon successful establishment, tumors were harvested and minced into 1-2 mm^3^ tumor bits and cryopreserved. Portions of the tumors were then further expanded subcutaneously in mice. All materials in this study are early passages.

#### Ex vivo culture establishment

The ex-vivo cultures were generated from PDXs by culturing in standard DMEM medium with 10% FBS or 10% Charcoal-Stripped Serum (CCS) and 5% Matrigel, without further additives. Briefly, the tumor was excised from subcutaneous xenografts around 500 mm^3^, rinsed with DMEM (gentamicin 50 μg/ml), and washed with DMEM and minced into 1-2 mm^3^ tumor bits. The tumor bits were digested with collagenase/dispase (Millipore Sigma, #10269638001 and #11097113001, respectively) at 37°C for 30 minutes. The digestion was then stopped by adding medium with 10% FBS, and the slurry went through 100 μM filter to remove big chunks. The tumor cells were collected by centrifugation. Resuspended cell pellets were then seeded in DMEM either with 10% FBS or 10% CCS and containing 5% (v/v) Matrigel (CORNING, catalog no. 356230), and plated at 0.8∼1 × 104 cells per well in 96-well plate.

#### CellTiter-Glo luminescent cell viability assay

Ex-vivo cultures generated from BIDPC4 PDX were cultured with DMEM plus either 10% FBS or 10%CCS and 5% Matrigel in 96-well plates. Cells were treated with nine doses of pan AKT inhibitors, serial diluting from 20 μM with a dilution factor of 2. Six replicates were set up for each dose. CellTiter-Glo luminescent cell viability assay (Promega, G7571) was performed after 7 days of treatment according to the manufacturer’s protocol. The killing curve was plotted based on the readout, which is normalized to the average luminescent signal of DMSO under the respective conditions. Two independent AKT inhibitors were used, MK2206 (MedchemExpress, Cat No. HY-10358) and ipatasertib (MedChemExpress, Cat /no. HY-15186). Experiments were repeated at least three times, and representative results were shown.

### Immunoblotting

Protein extracts were immunoblotted with antibodies against AKT pS473 (Cell Signaling rabbit mAb #4058, 1:1000) or pT308 (Cell Signaling rabbit mAb #2965, 1:1000), S6 pS235,236 (Cell Signaling #2211, 1:000), ERK pT202,Y204 (Cell Signaling rabbit mAb #4858, 1:1000), AR (mAb 441, Abcam #ab9474, 1:1000), ERG (Cell Signaling rabbit mAb #97249, 1:1000), Vinculin (Cell Signaling rabbit mAb #13901, 1:1000), PRAS40 (Cell Signaling rabbit mAb #13175, 1:1000), INPP4B (Cell Signaling rabbit mAb #14543, 1:1000), IRS2 (Cell Signaling rabbit mAb #4502, 1:1000), HA (Covance rabbit polyclonal #PRB-101C, 1:1000), AKT (Cell Signaling Technology #9272, 1:1000), AKT1 (Cell Signaling Technology #2938, 1:1000), AKT1 pS473 (Cell Signaling Technology #9018, 1:1000), AKT2 (Cell Signaling Technology #3063, 1:1000), AKT2 pS474 (Cell Signaling Technology #8599, 1:1000), PTEN (Cell Signaling Technology #9559, 1:1000), PHLPP1 (Abcam #ab305295, 1:1000), and PHLPP2 (Bethyl Laboratories #A300-661A, 1:2000).

### Immunohistochemistry

Primary antibodies were anti-pAKT (pS473, Cell Signaling rabbit mB #4058, 1:200), anti-HA (Cell Signaling rabbit mAb #3724, 1:200), anti-pS6 (S240/244, Cell Signaling, #2211, 1:200), anti-AR (Upstate rabbit polyclonal #06-680, 1:100), anti-TMPRSS2 (Cell Signaling rabbit mAb #84382), and anti-ERG (Abcam rabbit mAb EPR3864 #ab92513, 1:100) in 1% BSA. These were incubated overnight at 4°C, followed by biotinylated secondary antibody and streptavidin-HRP (1:400, Vector).

For NCT02903368 clinical trial specimens, we analyzed individual pretreatment core biopsies and a series of TMAs containing residual tumor from the post therapy radical prostatectomies. The TMAs contained 3 mm punches taken from 2-4 representative tumor foci based on histology. The *T:E* fusion status of the cases was determined by immunohistochemistry using an ERG antibody (2). Staining for pAKT was scored based on the fraction of positive cells, with a score of 0 for none, 1 for 1-5%, 2 for 5-25%, 3 for 25-50%, and 4 for >50%. The focus on the TMA from each case with the highest score was used for comparison with the pretreatment core biopsy. The scoring was carried out independently by two investigators who were blinded to ERG status.

For NCT02430480 clinical trial specimens, whole sections of biopsy and prostatectomy tissue stained with anti-ERG antibodies were scanned on a Carl Zeiss AxioScan.Z1 microscope slide scanner equipped with a Plan-Apochromat 20× NA 0.8 objective, 266% LED intensity, 200 µs exposure time. Tissue images were acquired using ZEN Blue 2012 (Zeiss) with objective/magnification and pixel:distance calibrations recorded within the scanned CZI file. Whole slide images were processed using HALO 3.4 (Indica Labs) with a Random Forest classifier for tumor detection and nuclear ERG quantification using the CytoNuclear detection module. All classified areas were reviewed by a pathologist and raw nuclear ERG intensity values (averaged per case) were report as optical densities.

### Transcriptome analyses

RNA from mouse prostate prior to and post DOX induction was analyzed by hybridization on Affymetrix Mouse Gene 1.0 ST arrays. For RNA-seq, mRNA libraries were generated using the Illumina TruSeq stranded mRNA sample kit. Raw reads were analyzed using a pipeline for RNA-seq analysis - Visualization Pipeline for RNA-seq analysis (VIPER) based on workflow management system Snakemake (https://github.com/hanfeisun/viper-rnaseq). The read alignment to the hg19 reference genome was performed using STAR aligner (2.7.0f) with default parameters. Gene expression (FPKM values) was quantitated with Cufflinks (v2.2.1). After ranking according to differential expression, Gene Set Enrichment Analysis (GSEA) was performed to search for enrichment across the Molecular Signatures Database (https://www.gsea-msigdb.org/gsea/index.jsp).

### ERG and gene set enrichment analyses of clinical samples

Whole-transcriptome sequencing data from matched pairs of tumors before and after neoadjuvant intense androgen deprivation therapy (GSE183100) were stratified by baseline anti-ERG IHC status (47). FASTQ pairs for GTEX prostate tissue data (phs000424.v10) were downloaded from Gen3. FASTQs for TCGA prostate cancer data (phs000178.v10) and the West Coast SU2C cohort (phs001648.v1) were downloaded from the NCI Genomic Data Commons. FASTQs for the East Coast SU2C cohort (phs000915.v2) were downloaded from dbGaP. TPM values were estimated using RSEM version 1.3.2 as a wrapper around STAR version 2.7.0f. Scaled estimates of gene expression at the TPM level in matched pairs of CRPC tumors from the West Coast SU2C cohort and Hartwig Medical Foundation (HMF) were previously published and provided by Xiaolin Zhu (39). Single-sample gene set enrichment scores were computed using raw or scaled TPM values using the GSVA package for R in ssgsea mode with tau = 0.75.

*T:E* fusion status from published RNA-seq cohorts lacking immunohistochemical annotation was determined using four complementary approaches. (1) If companion whole-genome sequencing data were available, structural variations involving *TMPRSS2* and *ERG* translocations or interstitial deletions were used to call a case as fusion-positive. (2) If companion whole-exome sequencing data were available, allele-specific imbalances between *TMPRSS2* and *ERG* (including inferred copy-number losses) were used to call a case as fusion positive. (3) In all cases, *de* novo assembly using defuse version 0.8.1 was employed to identify chimeric read-pairs mapping to the *T:E* mRNA. (4) Samples with outlier high levels of *ERG* expression lacking any other evidence of a fusion were manually inspected using the Integrative Genome Viewer to confirm it was truly *T:E* negative.

Gene expression correlations of prostate cancer TCGA data were performed directly in cBioPortal (48, 49).

### Chromatin immunoprecipitation sequencing analysis

Chromatin immunoprecipitation sequencing (ChIP-seq) data for ERG, H3K27Ac, H3K4me3, H3K4me1 were obtained from following list. ChIP-seq profiles of ERG and multiple histone modifiers flanking the IRS2 gene were demonstrated using IGV reference to Hg38.

| Sample name | GEO |
| --- | --- |
| VCaP NTC DHT *ERG* | GSM2086309 |
| VCaP SICTL H3K27AC CHIP-SEQ, REP2 | GSM2537220 |
| VCaP SI*ERG* H3K27AC CHIP-SEQ, REP2 | GSM2537223 |
| VCaP_EthI_H3K4me3 | GSM353603 |
| VCaP_regular_medium_H3K4me1 | GSM353631 |
| LNCaP_*ERG* | GSM353648 |
| H3K27AC LNCaP-DHT | GSM1249447 |
| LNCaP_regular_medium_H3K4me3 | GSM353626 |
| LNCaP_regular_medium_H3K4me1 | GSM353634 |

### Statistical analysis

Differences between two groups was determined by two-sided Student’s t tests and Wilcoxon rank sum tests. Both Pearson and Spearman correlation were used for correlation analysis. Statistical significance was accepted at P < 0.05. Nonlinear regression curve fitting with a variable slope model was applied to assess the dose-response of AKT inhibitors, and IC50 generated for each inhibitor under both FBS and CCS conditions. Statistical analyses were performed using GraphPad Prism 9 and RStudio.

### Study approval

The analysis of archival deidentified clinical samples was approved by the Beth Israel Deaconess Medical Center (BIDMC) IRB. All animal studies were approved by the BIDMC. Institutional Animal Care and Use Committee (IACUC) and conformed to the NIH guidelines.

## Data availability

Gene expression data generated in this study has been deposited in GEO with accession numbers GSE289099 and GSE289100. Any further data not included in the report can be obtained from the corresponding author.

## Author contributions

Conceptualization: FM, SC, SPB; Methodology: FM, SC, SA, LP, OV, ATK, CC, DJE, HY, AT, MET, AGS; Investigation: FM, SC, LC, BE, CC, SA, LC, FX, LP, OV, ATK, CC, DJE, FX, AG, ML, LP, OV, ATK, CC, HY, JWR, XY, MET, AGS; Visualization: FM, SC, LP, ATK, AGS; Funding acquisition: XY, MET, SPB; Project administration: SPB; Supervision: AGS, SPB; Writing-original draft: FM; Writing-editing: SC, SPB.

## Acknowledgments

This work was supported by NIH R01 CA168393 and CA272934-01 (SPB, XY), NIH P01 CA163227 (SPB), NIH P50 CA272390 (SPB, MET), NIH U54 CA156734 (SPB, CC), Department of Defense PCRP Idea Development Award W81XWH-20-1-0925 (FM, SPB), Department of Defense PCRP Idea Development Award W81XWH-15-1-0151 (XY), Prostate Cancer Foundation Challenge Awards (SPB, MET), a Research Fellowship from Gunma University Hospital (SA), and the Intramural Research Program of the National Cancer Institute (AGS). We thank Drs. John Clohessy and Johann de Bono for sharing data and helpful discussions.

---

**Figure S1.**
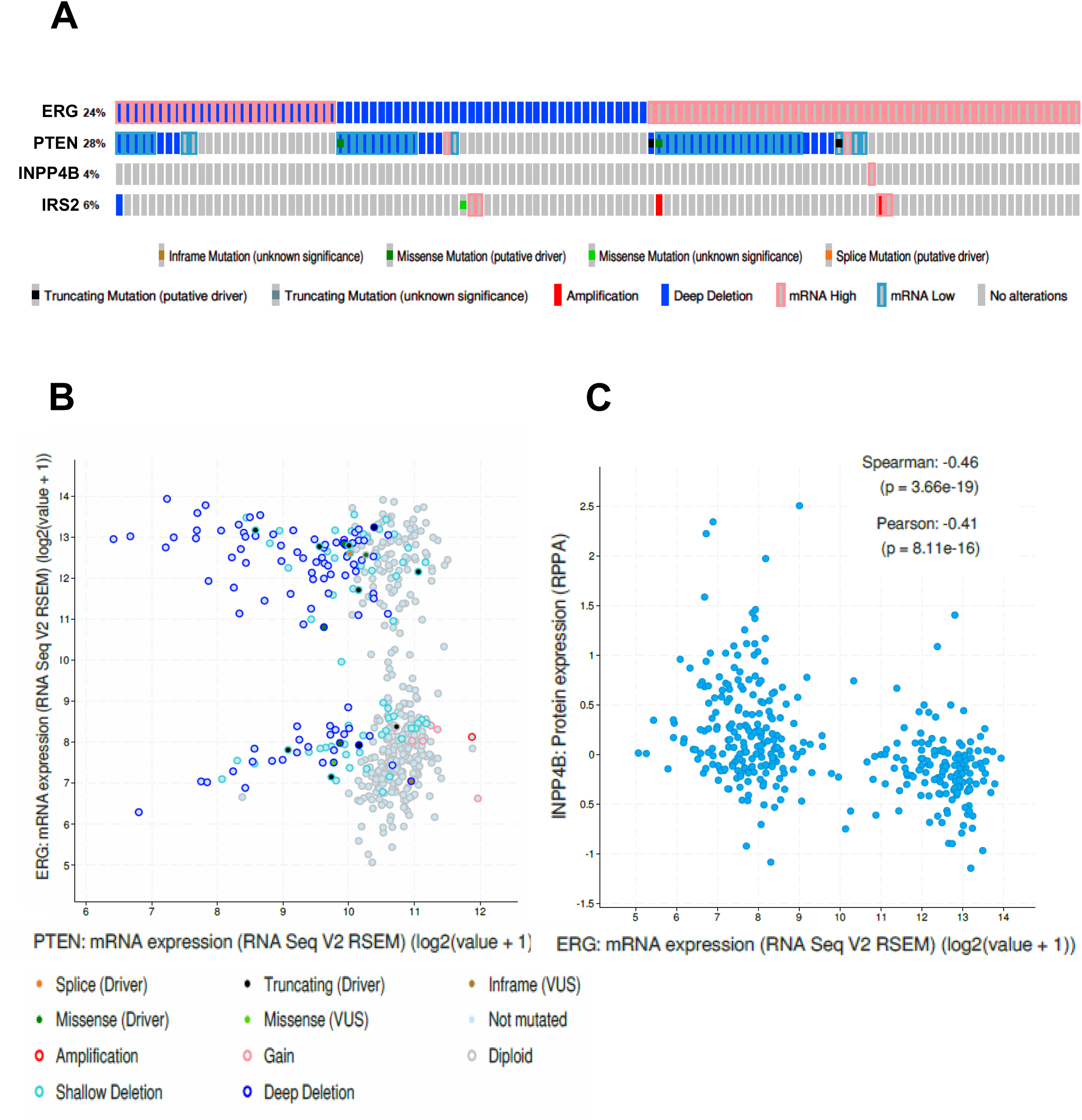
Associations between *T:E* fusion and PTEN in primary PCa. **(A)** Oncoprint showing alterations in *PTEN*, *INPP4B*, and *IRS2* in *T:E* fusion positive tumors in TCGA. Note that a subset of the *T:E* fusion positive tumors are scored as deep deletions, which likely reflects the fusion being mediated by interstitial deletion. **(B)** Correlation between ERG and PTEN mRNA in TCGA primary PCa showing PTEN mRNA levels in *PTEN* intact tumors are not decreased in *T:E* fusion tumors. **(C)** Correlation between ERG mRNA and INPP4B protein in TCGA primary PCa.

**Figure S2.**
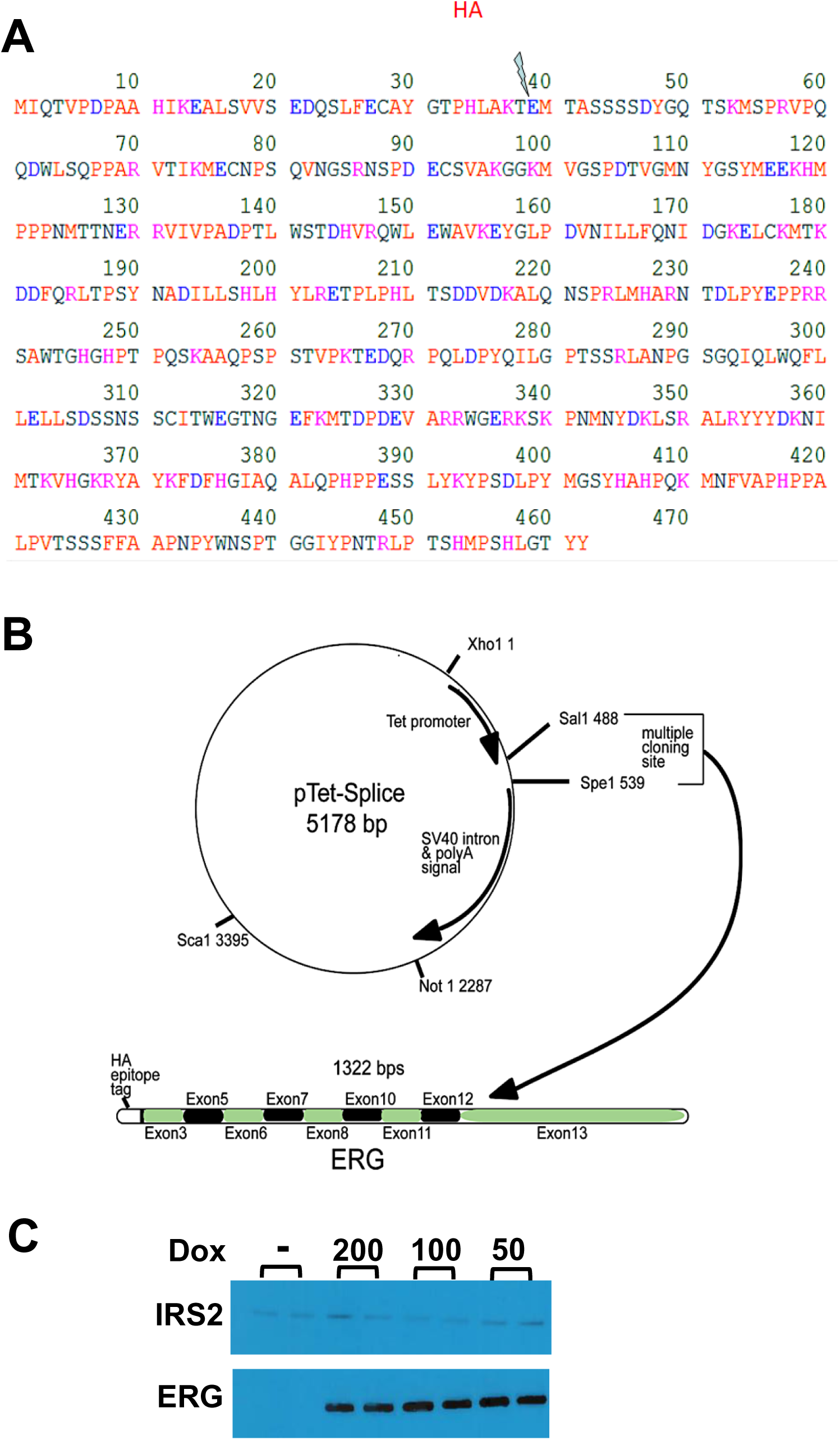
Construction of pTET-ERG expression vector. **(A)** HA epitope tag was inserted into human ERG with deletion of amino acids 1-39. **(B)** The HA-tagged ERG was cloned into the pTET-Splice plasmid to yield pTET-ERG. **(C)** LNCaP cells stably expressing the DOX-inducible HA-tagged ERG were treated with DOX (0 -200 ng/ml) for 2 days and assessed by immunoblotting for IRS2 and ERG.

**Figure S3.**
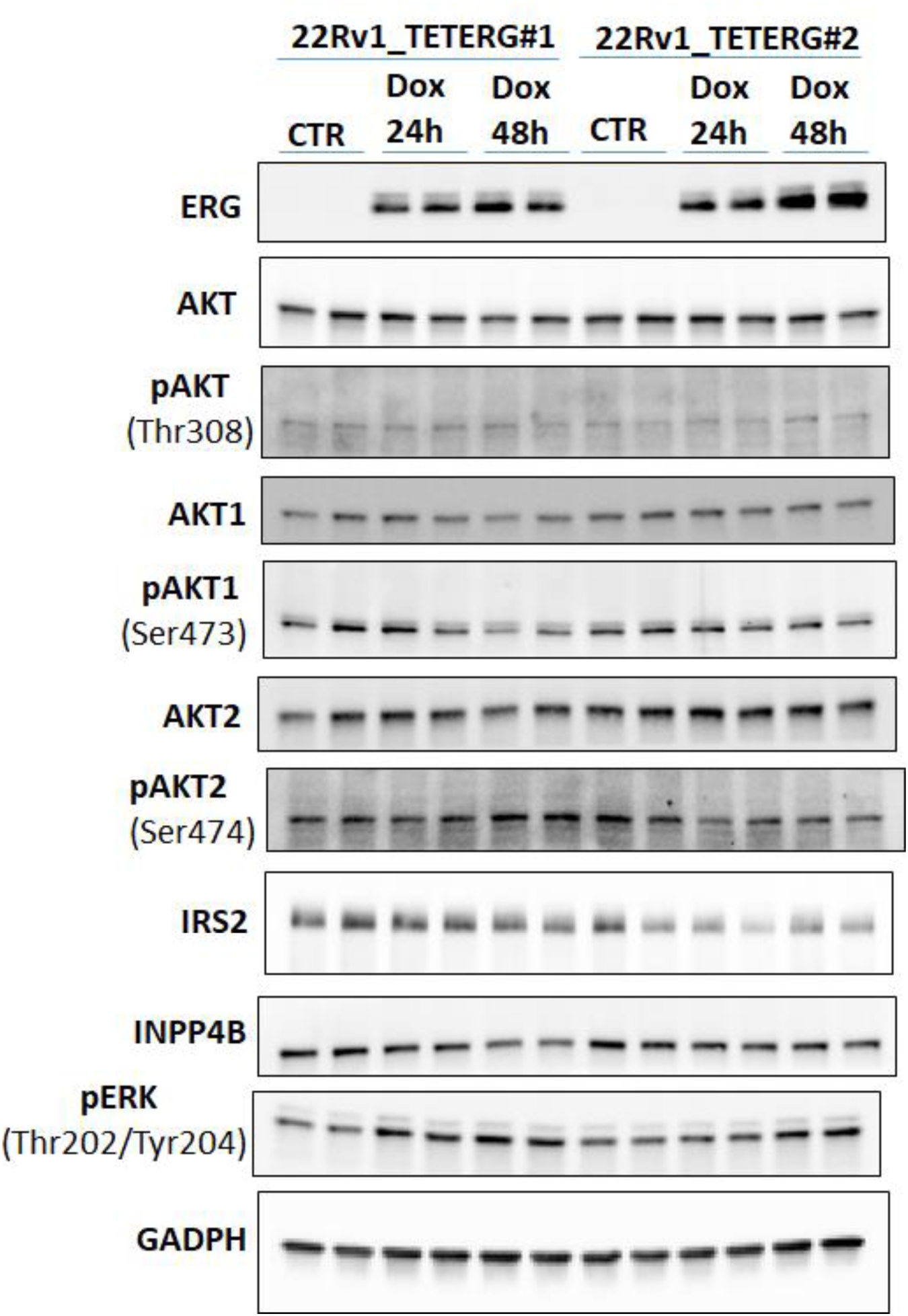
Effects of acute ERG induction in 22Rv1 cells. Two independent 22Rv1 lines with Dox-inducible expression of ERG were treated with Dox (50 ng/ml) versus vehicle for 48 hours followed by immunoblotting as indicated. Biological replicates are shown for each condition.

**Figure S4.**
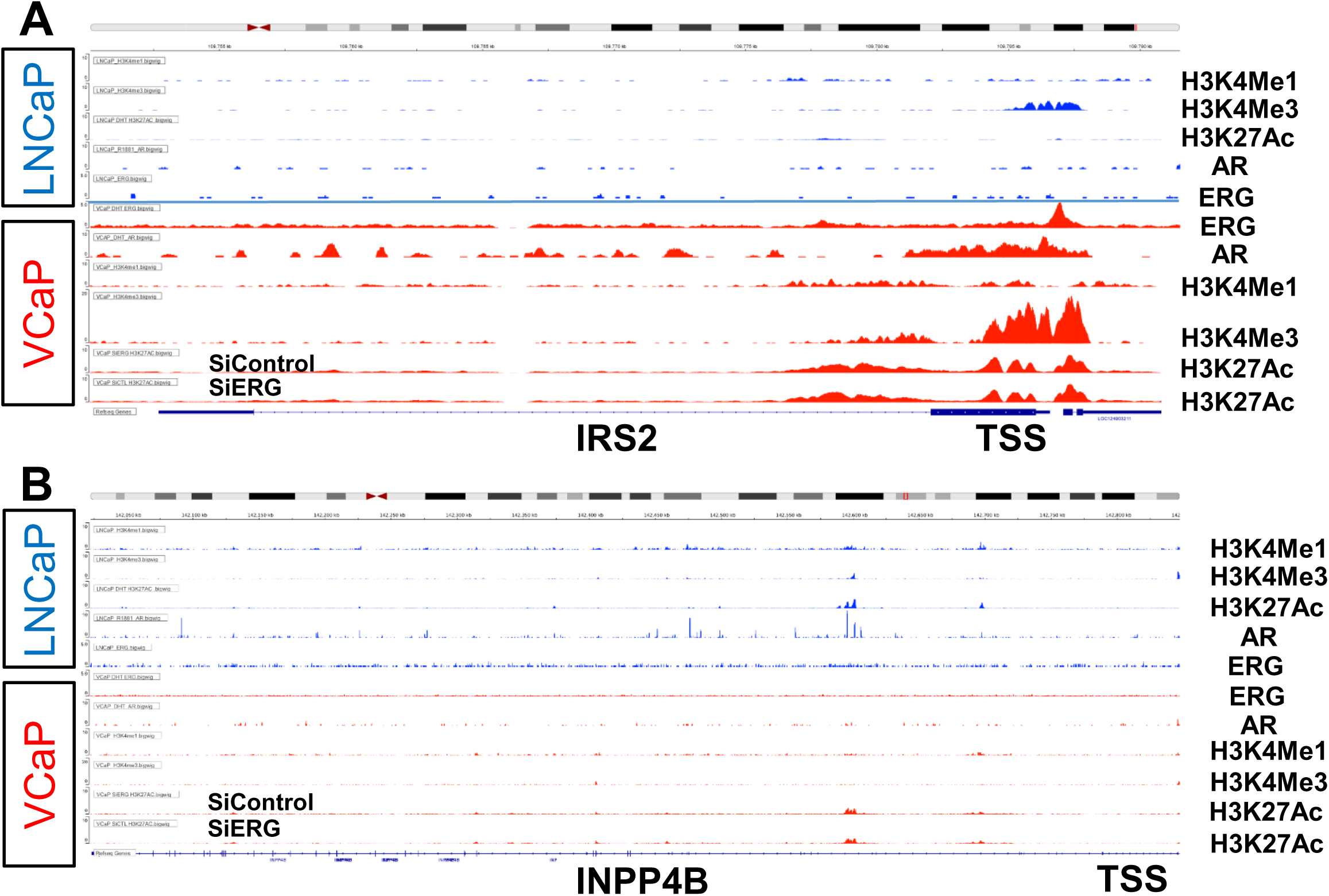
Binding profile of AR and ERG to the IRS2 and INPP4B transcription start sites in VCaP (*T:E* fusion positive) and LNCaP (*T:E* fusion negative) PCa cells. **(A)** ChIP-seq profiles for ERG, AR, and histone modifications H3K27Ac, H3K4Me3, and H3K4Me1 across a segment of genomic DNA encompassing *IRS2*. Each row represents a different ChIP-seq experiment with peaks indicating regions of protein-DNA interaction or histone modification enrichment. All data were retrieved from GEO, with accession number listed in Methods. Data ranges were set as 0-5 for ERG track, 0-20 for H3K4Me3 VCaP track, and 0-10 for all other tracks. **(B)** Binding profiles at *INPP4B* gene. Data ranges were set as 0-5 for ERG track, and 0-10 for all other tracks.

**Figure S5.**
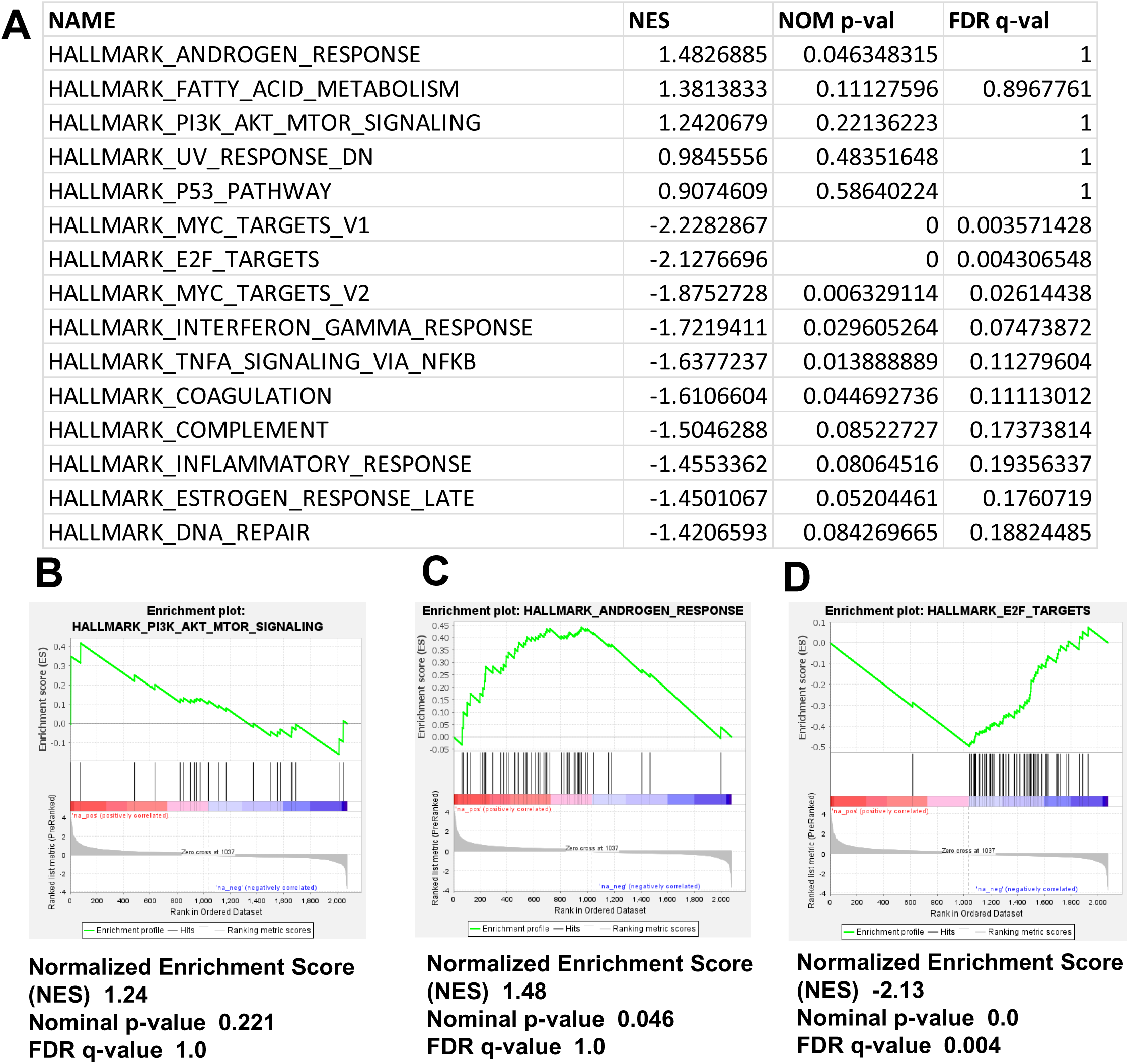
Gene set enrichments in siERG versus control VCaP cells. **(A)** Gene sets most highly increased and decreased in VCaP cells treated with nontarget control siRNA versus ERG targeted siRNA. Gene sets were determined using the GSEA browser (www.gsea-msigdb.org) version 4.4.3 and a rank list with genes that were significantly altered (p <.05) is shown. **(B-D)** Enrichment for Hallmark PI3K AKT MTOR (B), Androgen Response (C), and E2F Targets (D) gene sets.

**Figure S6.**
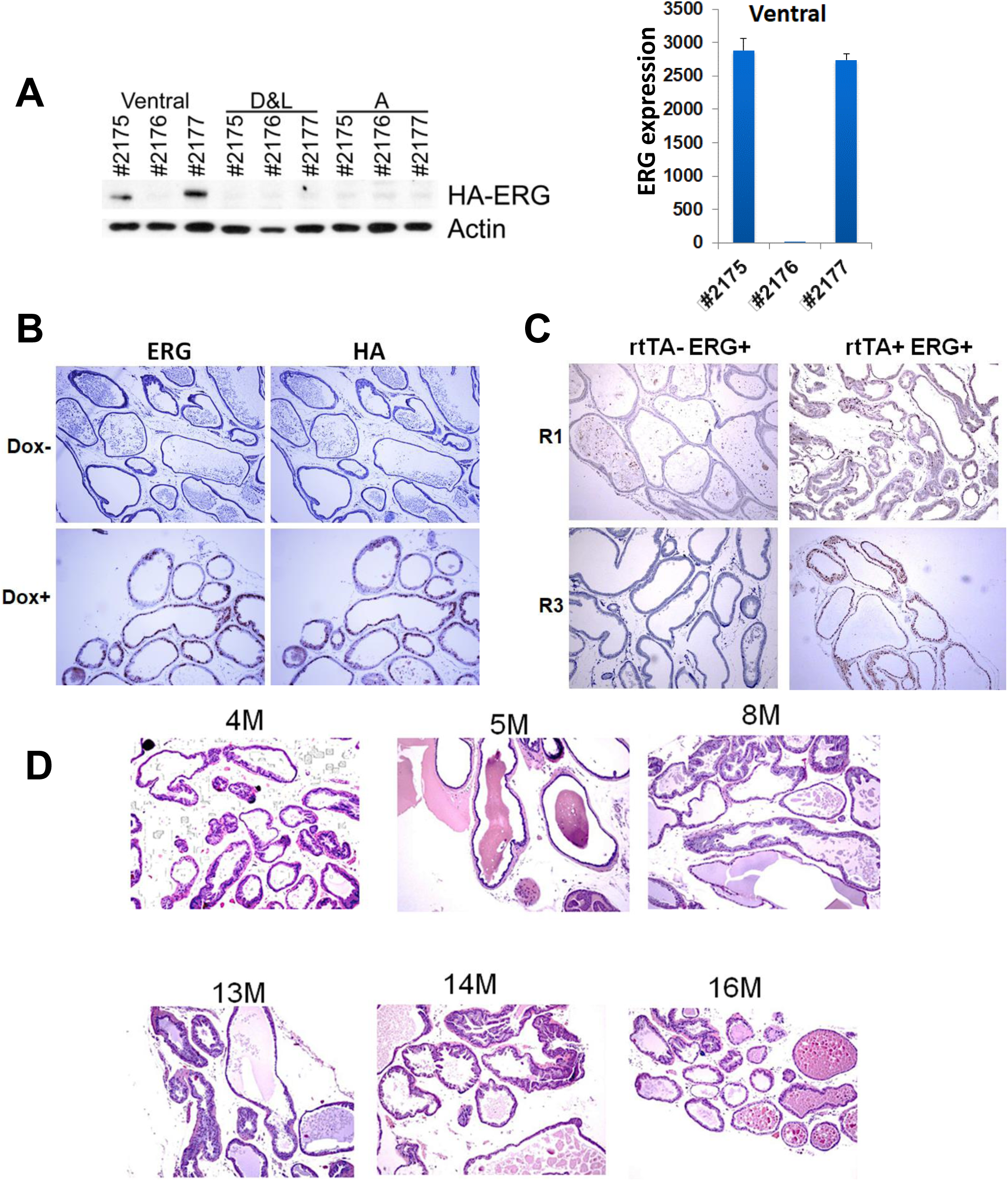
Doxycycline induction of ERG in mouse prostate. **(A)** Mice expressing probasin-rtTA and tet0-HA-ERG were fed doxycycline chow for ∼4 weeks and then sacrificed. Prostates were dissected into ventral, dorsolateral (D&L), and anterior (A) lobes and assessed for ERG by immunoblotting (left). Ventral lobe was further assessed by qRT-PCR for human ERG, normalized to untreated prostate (right). **(B)** ERG and HA expression in ventral prostate after 24 days of DOX or control chow. **(C)** Two independent lines of tetO-ERG mice (R1 and R3) were bred to coexpress the probasin-rtTA transgene (rtTA+ ERG+) in comparison with those not coexpressing probasin-rtTA (rtTA- ERG+). Mice were fed DOX chow and sacrificed to assess ERG induction, which confirmed the ERG expression was probasin-rtTA dependent. **(D)** Representative images of prostates from probasin-rtTA; tetO-ERG mice fed DOX chow for 4 – 16 months.

**Figure S7.**
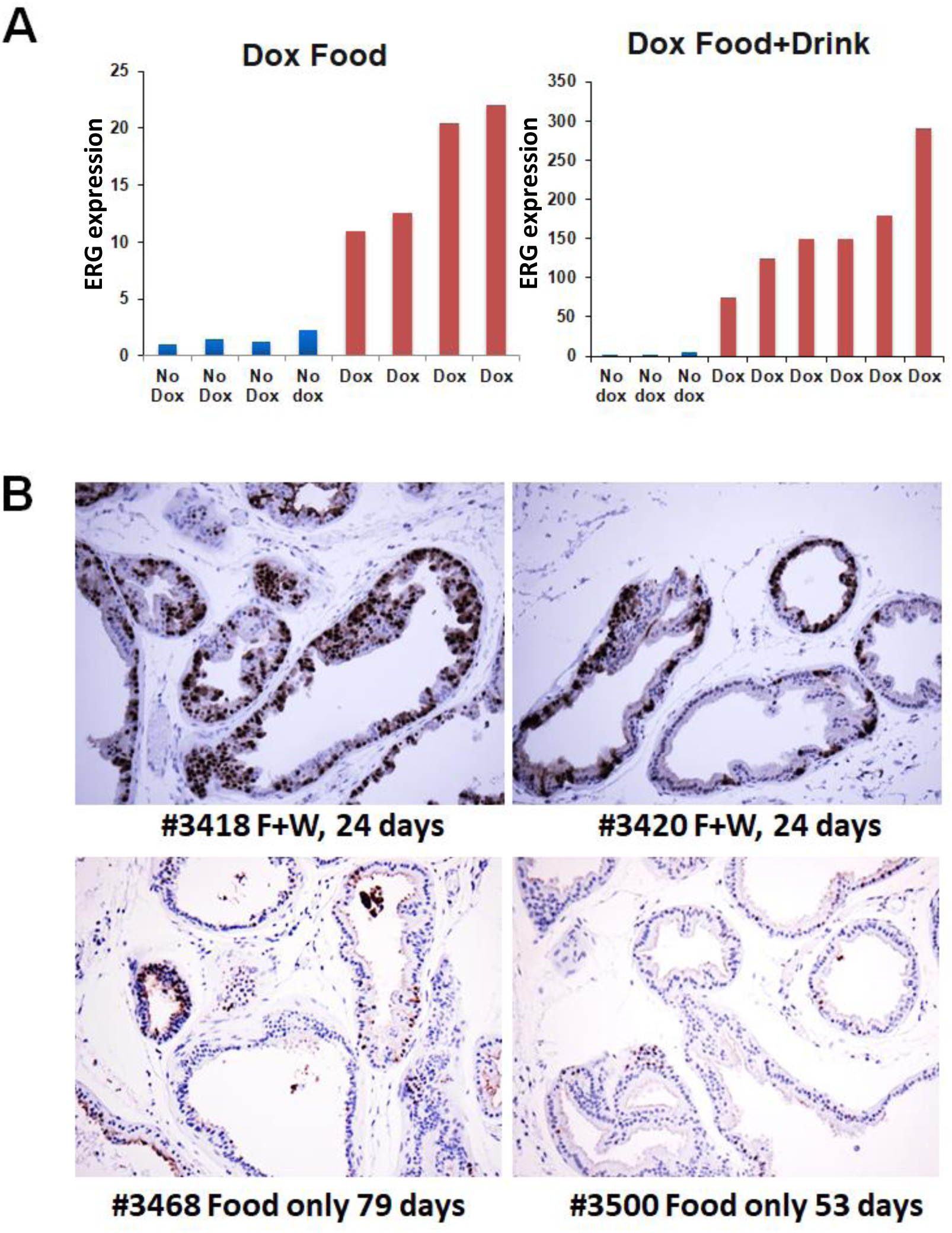
ERG expression is increased by adding doxycycline to water in addition to chow. Probasin-rtTA;tetO-ERG;*Pten*^+/-^ mice were treated with DOX in chow or in chow plus water (or no DOX) for ∼1-3 months. ERG expression in ventral prostate was then assessed by qRT-PCR (normalized to no Dox) **(A)** or IHC **(B)**.

**Figure S8.**
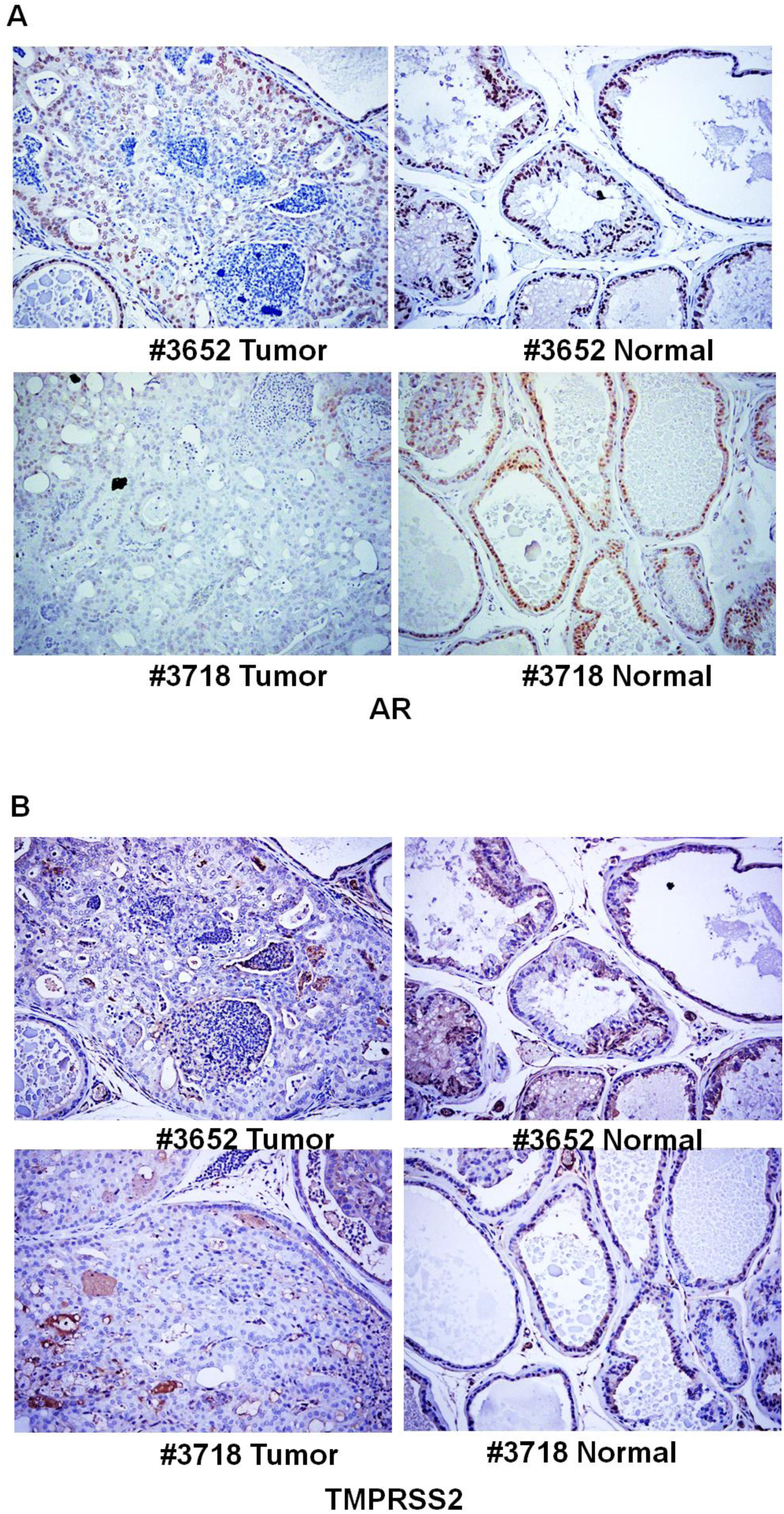
Decreased expression of AR and TMPRSS2 in tumor foci. **(A)** Probasin-rtTA;tetO-ERG;*Pten*^+/-^mice treated with DOX (food and water) for ∼12 months were sacrificed. Areas showing tumor and normal appearing areas were identified by H&E and then assessed by IHC for AR. **(B)** Additional sections from blocks in (A) were stained for TMPRSS2.

**Figure S9.**
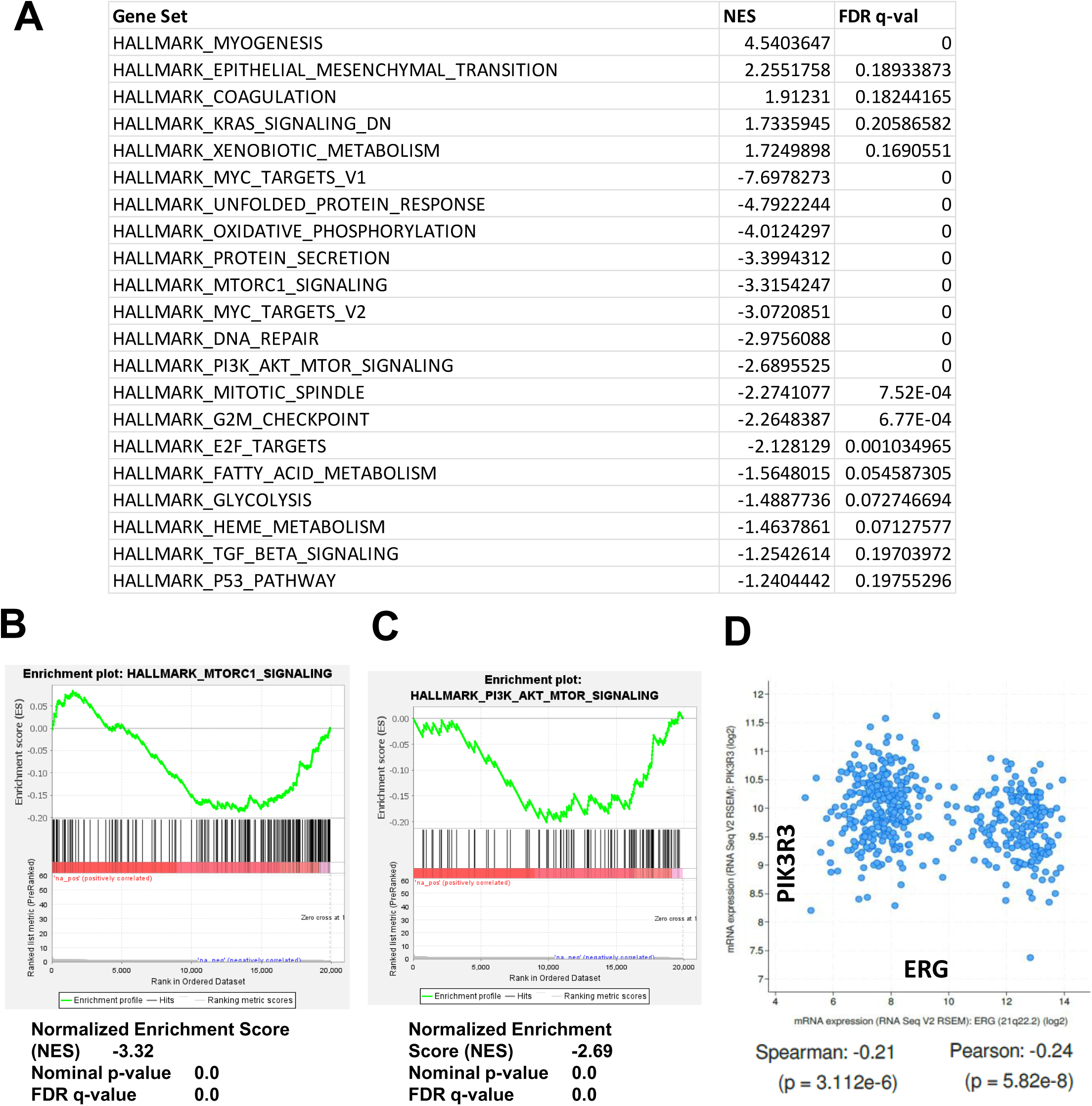
Hallmark gene set enrichments in response to initiating doxycycline in probasin-rtTA;tetO-ERG;*Pten*^+/-^ mice. **(A)** Ventral prostate was isolated from mice treated for 3-6 days with DOX (food and drinking water) or control. RNA was extracted and analyzed on Affymetrix microarrays, followed by GSEA. Hallmark gene sets altered (increased or decreased) with FDR <0.25 are shown. **(B)** Enrichment for MTORC1 Signaling in response to DOX induction. **(C)** Enrichment for PI3K AKT MTOR Signaling in response to DOX induction. **(D)** Correlation between *T:E* fusion and PIK3R3 in TCGA primary PCa data set.

**Figure S10.**
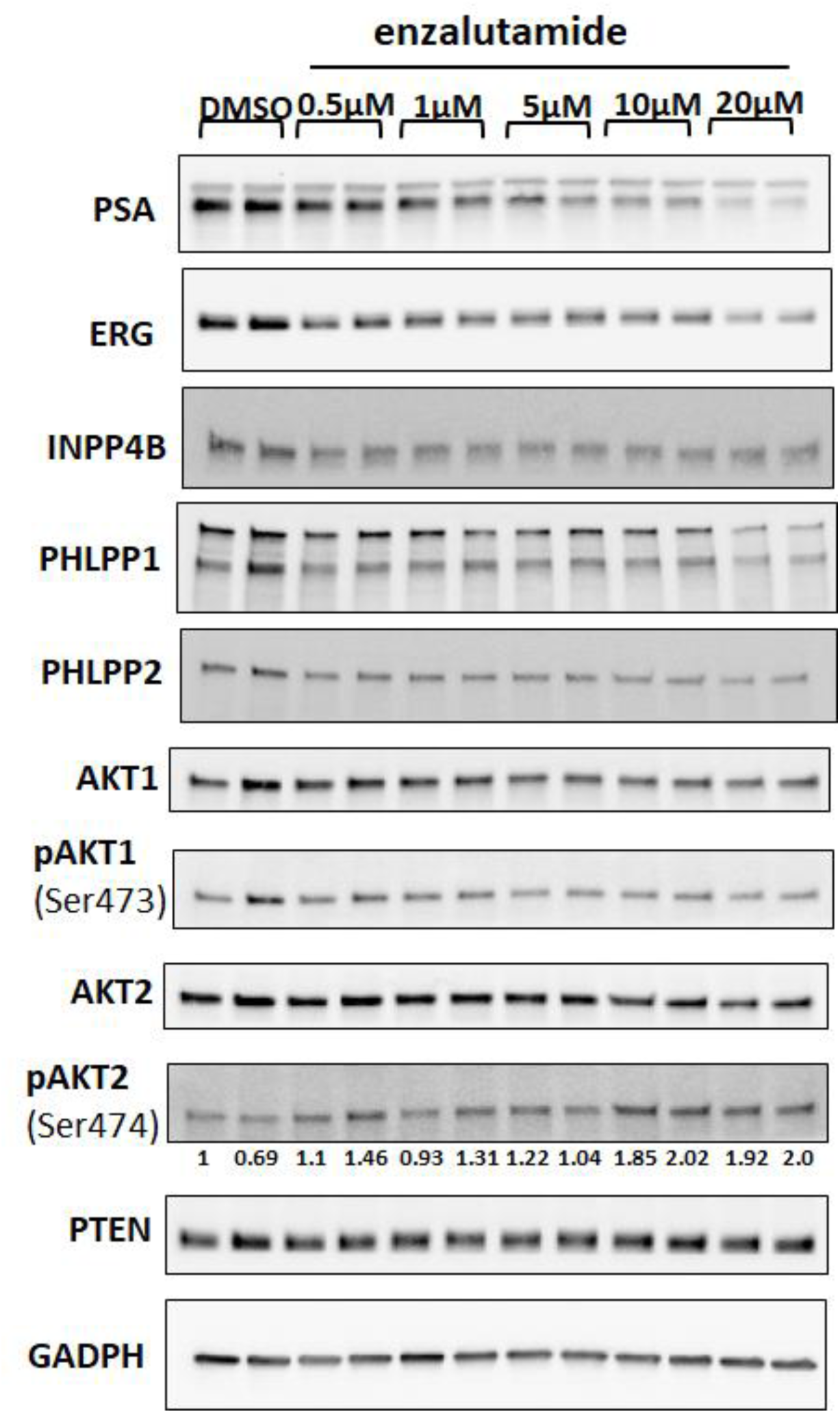
Effects of AR inhibition of ERG and AKT activation in *T:E* fusion cells. VCaP cells were treated for 48 hours with enzalutamide (0 – 20 μM) followed by immunoblotting as indicated. Biological replicates are shown for each condition. Band intensities for pAKT2 were quantified and values shown are normalized to the level in the first control.

**Figure S11.**
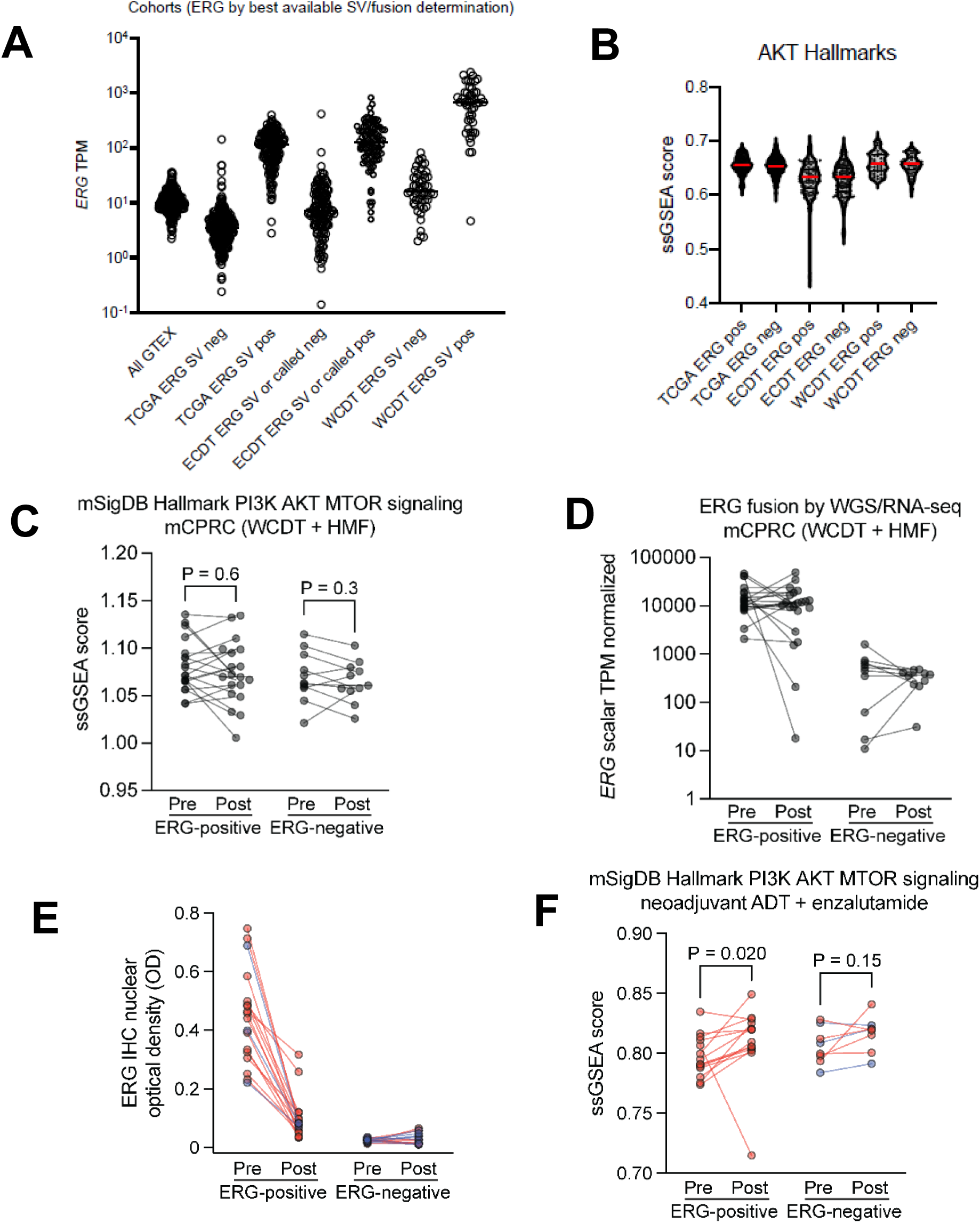
AKT activity is not increased in *T:E* positive CRPC. **(A)** Samples were separated into *T:E* fusion positive versus negative tumors based on genomic determination of structural variation/fusion status and each group was then assessed for ERG expression (transcripts per million, TPM) based on RNA-seq data. Samples are nonneoplastic prostate (GTEX, n= 243), TCGA primary PCa (n = 500), East Coast Dream Team CRPC (ECDT, n = 236), West Coast Dream Team (WCDT, n = 99). (B) ssGSEA showing enrichment for the Hallmark PI3K_AKT_MTOR gene set in each tumor. (**C, D**) Hallmark PI3K_AKT_MTOR gene set enrichment (**C**) and ERG expression (**D**) in matched CRPC tumors before and after AR-targeted therapy. (**E, F**) Tumors from figure 6 that are PTEN intact are shown in blue. The ssGSEA score in *PTEN* intact *T:E* tumors and some *T:E* negative tumors could not be determined due to low levels of residual tumor for which RNA-seq could not be done.

**Figure S12.**
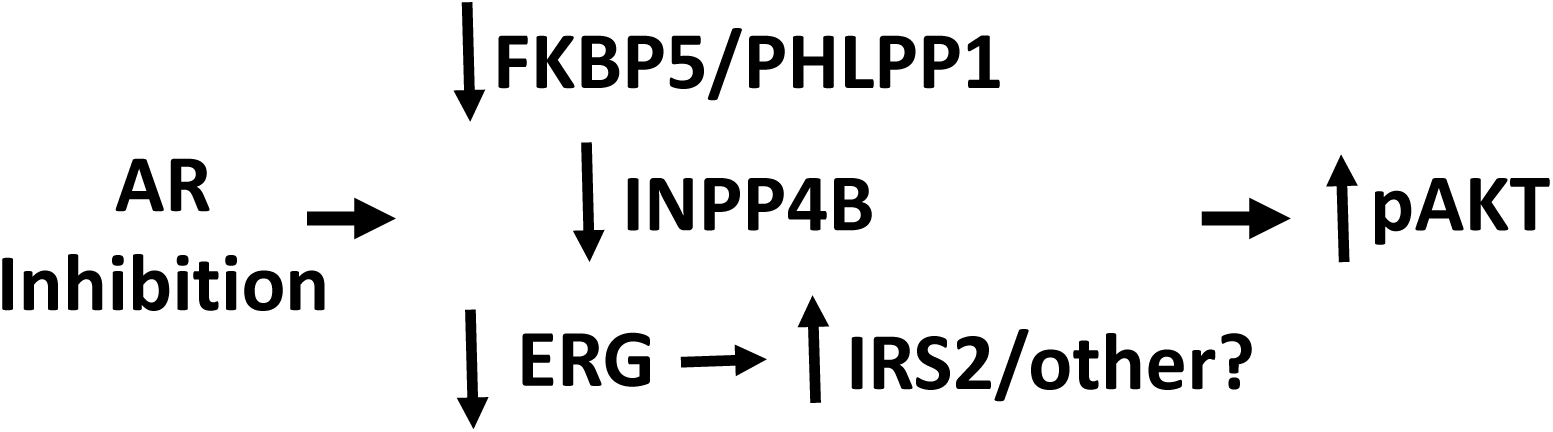
Potential mechanisms driving AKT activation in response to AR inhibition in *T:E* fusion tumors. AR inhibition decreases expression of FKBP5, which may act as a scaffold for PHLPP1-mediated dephosphorylation of AKT, leading to increased pAKT. AR inhibition also decreases INPP4B, leading to increased levels of PI(3,4)P_2_ that can mediate AKT recruitment and activation. AR inhibition in *T:E* fusion tumors also decreases ERG, which can increase IRS2 and thereby enhance signaling by receptor tyrosine kinases and downstream activation of PI3K/AKT, and potentially by other pathways. The decrease in ERG may also act through other mechanisms that remain to be determined. Notably, the decrease in INPP4B may further amplify this increase in receptor tyrosine kinase signaling.

